# The β-cell primary cilium is an autonomous Ca^2+^ compartment for paracrine GABA signalling

**DOI:** 10.1101/2021.08.16.456473

**Authors:** Gonzalo Sanchez, Tugce Ceren Incedal, Juan Prada, Paul O’Callaghan, Santiago Echeverry, Phuoc My Nguyen, Beichen Xie, Sebastian Barg, Johan Kreuger, Thomas Dandekar, Olof Idevall-Hagren

## Abstract

The primary cilium is an organelle present in most adult mammalian cells and is thought of as an antenna for detection of a variety of signals. Here we use intact mouse pancreatic islets of Langerhans to investigate signalling properties of the primary cilium in β-cells. Using cilia-targeted Ca^2+^ indicators we find that the resting Ca^2+^ concentration in the cilium is lower than that of the cytosol, and we uncover a Ca^2+^ extrusion mechanism in the cilium that effectively insulates the cilium from changes in cytosolic Ca^2+^. Stimuli that give rise to pronounced cytosolic Ca^2+^ concentration increases, such as glucose- and depolarization-induced Ca^2+^ influx, and mobilization of Ca^2+^ from the ER, was accompanied by minor increases in cilia Ca^2+^ concentrations that were spatially restricted to a small compartment at the base. Conversely, we observe pronounced Ca^2+^ concentration changes in the primary cilia of islet β-cells that do not propagate into the cytosol and show that paracrine GABA signalling via cilia-localized GABA-B1-receptors is responsible for this Ca^2+^ signalling. Finally, we demonstrate that the cilia response to GABA involves ligand-dependent transport of GABA-B1 receptors into the cilium.

## INTRODUCTION

The primary cilium is a specialized signalling organelle interfacing the cell and its environment and with the capability of translating multimodal inputs and prompting appropriate cellular responses. The presence of specific ciliary receptors and effector proteins enables the generation of local ciliary signals and provides a platform for selective crosstalk between different pathways, with Ca^2+^ and cAMP being the most prominent second messengers in this organelle ^1–4^ Ca^2+^ in particular has been associated with different ciliary responses but how ciliary Ca^2+^ signalling is protected from cytoplasmic interference and what events take place upstream of the second messenger are not well understood. The concentration of Ca^2+^ in the cilium has been reported to be higher than in the cytosol due to constitutive influx ^5^. However, such an arrangement is not readily reconcilable with traditional Ca^2+^ signalling downstream of cell surface receptors, and it would also greatly limit the number of Ca^2+^ effectors that can be used to decode the Ca^2+^ signals, since many have affinities for Ca^2+^ in the nanomolar range. Ca^2+^ has also been proposed to freely diffuse between the cilium and cytosol ^5,6^, an arrangement that is difficult to reconcile with a specific role of Ca^2+^ in the cilium, since that would require a mechanism for distinguishing Ca^2+^ of cytosolic and ciliary origin. Many G-protein coupled receptors (GPCRs) localize to primary cilia of a variety of cell types and the current model for signal modulation postulates that GPCRs are removed from the cilia upon agonist activation for signal termination ^7^. However, most of the details concerning ciliary signalling downstream of receptor activation remain elusive. In vertebrates, Hedgehog signaling relies on the cilium and, despite the complexity of the Hedgehog pathway, it is perhaps the best-known example of ciliary signal transduction. The Hedgehog receptor Patched is mainly localized to the ciliary membrane and its activation results in exit from the cilium, allowing entry of Smoothened and subsequent activation of the pathway ^8^.

Pancreatic β-cells are the only source of insulin, a major anabolic hormone important for maintaining blood glucose homeostasis. Failure to produce or secrete insulin results in elevated blood glucose concentrations and contribute to the development of diabetes ^9^. β-cells are located within islets of Langerhans, which are richly vascularized endocrine cell clusters dispersed throughout the pancreas. Each β-cell is equipped with a primary cilium that protrudes from the apical side facing away from the islet vasculature ^10,11^. β-cell-specific loss of primary cilia is associated with impaired cell function, but the signaling downstream of this organelle is still poorly understood ^12–14^. In particular, a role of the β-cell cilia as an antenna sensing the islet microenvironment has not been demonstrated.

GABA is mainly regarded as the inhibitory neurotransmitter in the brain even though it is an evolutionary ancient signaling molecule and a variety of other actions outside of the central nervous system have been documented ^15^. In the endocrine pancreas, for instance, GABA has both acute and long-term effects, including modulation of insulin secretion and maintenance of β-cell identity and function ^16,17^. β-cells produce and release GABA that acts locally within the pancreatic islets of Langerhans, though the mechanisms behind its signaling are still not well understood ^18^.

In the present work, we used intact endocrine pancreatic islets of Langerhans to measure ciliary Ca^2+^ dynamics within these functional micro-organs. Contrary to the vast majority of studies on the primary cilium in immortalized cell lines, the current work enabled the study of cilia signaling in a preparation that preserves both tissue architecture and function. Using this model, we unveiled cilia-specific Ca^2+^ activity driven, at least in part, by activation of metabotropic GABA receptors and that depends on Ca^2+^ extrusion, which isolates the cilium from contaminating cytosolic Ca^2+^. We describe here for the first time a cilia-localized class C GPCR, the GABA-B1 receptor, and show that it exhibits a distinct distribution with enrichment towards the cilia base. We also show that agonist stimulation mobilizes receptors to more distal parts of the cilium and triggers ciliary Ca^2+^ transients in a putative non-canonical fashion.

## MATERIALS AND METHODS

### Plasmids, adenoviruses and reagents

The following plasmids were used: Lyn_11_-R-GECO^19^, 5HT_6_-G-GECO (Addgene plasmid 47499; kind gift from Takanari Inoue^6^), 5HT6-Venus-CFP (Addgene plasmid 47501; kind gift from Takanari Inoue^6^), ss-pHluorin-Smo (Kind gift from Derek Toomre, Yale Univeristy^20^). Smo-GCaMP5G-mCherry was generated by PCR-amplification of GCaMP5G (Addgene plasmid 31788, kind gift from Loren Looger^21^) using the following primers: GCaMP-EcoRI-Fwd (tatata<u>GAATTC</u>TAatgggttctcatcatca) and GCaMP-SacII-Rev (ctctct<u>CCGCGG</u>cttcgctgtcatcatt) followed by digestion of the PCR product with EcoRI and SacII and ligation in between Smoothened and mCherry in mCherry-Smo (Addgene plasmid 55134, kind gift from Michael Davidson). Adenoviral particles (E5 serotype) carrying 5HT_6_-G-GECO and Smo-GCaMP5G-mCh under the control of CMV-promoters were produced by Vector Biolabs (Malvern, PA). All salts, Mg-ATP and α-toxin were from Sigma-Aldrich/Merck (Kenilworth, New Jersey, USA). GABA, diazoxide, methoxyverapamil, α-ketoisocaproic acid, nifedipine, forskolin, baclofen, CGP35348, picorotoxin, amiloride, ATP, carbachol and vigabatrin were from Tocris (Bristol, UK). The following antibodies were used in the study: acetylated tubulin (T7451 Sigma, host: mouse, 1:500), IFT88 (F41236 NSJ Bioreagents, host: rabbit, 1:200), SSTR3 (E-AB-1607 Elabscience, host: rabbit, 1:200), Patched (ab53715 Abcam, host: rabbit, 1:200), Smoothened (ab113438 Abcam, host: rabbit, 1:200), GABA-B1 receptor (NBP1-52389 Novus Biologicals, host: goat, 1:200. And AGB-001Alamone Labs, host: rabbit, 1:200), GABA-B2 receptor (ab230136 Abcam, host: rabbit, 1:200), GABA-A receptor (224-103 Synaptic Systems, host: rabbit, 1:200). Secondary antibodies for confocal microscopy were anti-mouse 568nm (A11004 Invitrogen) and anti-mouse 488nm (A28175 Invitrogen), anti-rabbit 488nm (A11034 Invitrogen), anti-goat 555nm (A21432 Invitrogen); all secondary antibodies were diluted to 1:500. For super-resolution microscopy the following secondary antibodies were used: anti-mouse Abberior Star 580 and anti-rabbit Abberior Star Red (Abberior GmbH, Göttingen, Germany). All secondary antibodies were diluted to 1:300.

### Cell- and islet culture and transfection

#### MIN6 cell culture and transfection with plasmids and siRNA

The mouse β-cell line MIN6 (passages 18-30)^22^ was cultured in DMEM (Life Technologies) supplemented with 25 mmol/l glucose, 15% FBS, 2 mmol/l L-glutamine, 50 μmol/l 2-mercaptoethanol, 100 U/ml penicillin and 100 μg/ml streptomycin. The cells were kept at 37°C and 5% CO_2_ in a humidified incubator. Prior to imaging, 0.2 million cells were resuspended in 100 μl Opti-MEM-I medium (Life technologies) with 0.2 μg plasmid (total) and 0.5 μl lipofectamine 2000 (Life technologies) and seeded in the centre of a 25-mm poly-L-lysine-coated coverslip. The transfection reaction was terminated after 4-6h by the addition of 2 mL complete culture medium and cells were imaged 18-24 h later. For knockdown experiments MIN6 cells were resuspended with 25 nM antiGABA-B1 receptor or control siRNA (Dharmacon, siGENOME) pre-mixed with 1.6 μl/ml Lipofectamine RNAiMAX (Life technologies) in Opti-MEM I medium. After 3 hours, the Opti-MEM I medium was replaced by DMEM growth medium containing 25 nM siRNA and 1.6 μl/ml Lipofectamine 3000 (Life technologies) and incubated overnight.

#### MIN6 pseudoislet formation and culture

Detached MIN6 cells (3 to 5 million/ml) were transferred to a non-adherent petri dish (Sarstedt, Nümbrecht, Germany) with 5 ml culture medium and kept in culture for 5-7 days at 37 °C in an humidified atmosphere with 5% CO_2_ until they spontaneously formed cellular aggregates (pseudoislets).

#### Mouse islets isolation and culture

Adult C57Bl6J mice (>8 months) were sacrificed by CO_2_ asphyxiation and decollation and the pancreas was removed and put on ice prior to digestion with collagenase P and mechanical disaggregation at 37°C. The digestion was terminated by 1 ml BSA 0.1 g/ml and islets were then separated from exocrine tissue by handpicking with a 10 μl pipette under a stereo microscope. Islets were cultured in RPMI 1640 medium with 5.5 mM glucose, supplemented with 10 % fetal calf serum, 100 units/ml penicillin, 100 μg/ml streptomycin for 2 to 5 days at 37 °C in a humidified atmosphere with 5% CO_2_. All procedures for animal handling and islet isolation were approved by the Uppsala animal ethics committee.

#### Human islet culture

Human islets from normoglycemic cadaveric organ donors were kindly provided via the Nordic Network for Clinical Islet Transplantation. All experiments with human islets were approved by the Uppsala human ethics committee. Islets were cultured for up to 5 days in CMRL 1066 medium containing 5.5 mM glucose and supplemented with 10% fetal calf serum, 100 units/ml penicillin and 100 μg/ml streptomycin and kept at 37 °C in an atmosphere of 5% CO_2_ in humidified air.

#### Viral transduction of islets and pseudoislets

Islets were transferred to a petri dish containing a 200 μl drop of culture medium and 10 of high titration virus (> 10^12^-10^13^ vp/ml; Vector Biolabs, Malvern, PA). After 3 h, islets were transferred to a new petri dish filled with 5 ml of fresh medium and kept in culture for at least 3 days to allow for biosensor expression before performing experiments.

### α-toxin permeabilization and preparation of intracellular-like media

Intracellular-like media with buffered pH, [Ca^2+^] and [Mg^2+^] used in α-toxin permeabilization experiments contained: 6 mM Na^+^, 140 mM K^+^, 1 mM (free) Mg^2+^, 0-100 μM (free) Ca^2+^, 1 mM Mg-ATP, 10 mM HEPES, 2 mM (total) EGTA and 2 mM (total) Nitrilotriacetic acid (NTA) with pH adjusted to 7.00 at 22°C with 2M KOH. The total concentration of Ca^2+^ and Mg^2+^ was calculated using the online version of MaxChelator (http://www.stanford.edu/~cpatton/webmaxcS.htm). Media were made fresh on the day of experiment and kept on ice. For permeabilization, 25-mm poly-L-lysine-coated glass coverslips with transfected, adherent MIN6 cells were used as exchangeable bottoms in a modified Sykes-Moore open superfusion chamber that was mounted on the stage of a TIRF microscope (described below) and connected to a peristaltic pump that allowed rapid change of medium. Following change from normal, extracellular-like, medium (125 mM NaCl, 4.9 mM KCl, 1.3 mM MgCl_2_, 1.2 mM CaCl_2_, 25 mM HEPES, 1 mg/ml BSA with pH set to 7.4) to an intracellular-like medium (see below), the superfusion was interrupted and alpha-toxin was added directly to the chamber (final concentration ≈50 μg/mL). Permeabilization typically took 2-5 min after which superfusion was started again and the cells were exposed to intracellular-like buffers containing calibrated Ca^2+^ concentrations while fluorescence from GCaMP5G and mCherry was recorded. These experiments were performed at ambient temperature (21-23 °C).

### Fluorescence microscopy

#### Confocal microscopy

A spinning disk confocal microscope unit (Yokogawa CSU-10) mounted on an Eclipse Ti2 body (Nikon, Japan) was used to acquire pictures of immuno-labelled samples. The microscope was equipped with a CFI Apochromat TIRF 100X, 1.49 NA oil immersion objective from Nikon (Japan) and excitation light was provided by 491 nm and 561 nm DPSS lasers from Cobolt (Hübner Photonics, Solna, Sweden). Lasers were merged with dichroic mirrors, homogenized using a rotating, light-shaping diffusor and delivered to the CSU. Excitation light source was selected by electronic shutters (SmartShutter, Sutter Instruments) and emission light was separated using the following filters fitted into a filter wheel controlled by a Lambda 10-3 unit (Sutter Instruments): GFP/Alexa488 (530/50, Semrock), mCherry/Alexa561 (593LP, Semrock). Images were captured with a back-illuminated EMCCD camera (DU-888, Andor technology, UK) using MetaFluor software (Molecular Devices, San Jose, CA). For the mechanical stimulation experiments shown in figure 3, alternative confocal microscope setups were used. For details, see section “Mechanical stimulation of islets and cells” below.

#### Total Internal Reflection Fluorescence (TIRF) microscopy

TIRF imaging was performed on a Nikon TiE microscope equipped with an iLAS2 TIRF illuminator for multi-angle patterned illumination (Gataca systems, France) and a 100X1.49-NA Apo-TIRF objective. Excitation light for GFP and mCherry was delivered by 488-nm and 561-nm diode-pumped solid-state lasers with built-in acousto-optical modulators, and light for bleaching or photorelease of Ca^2+^ was delivered by a 405-nm DPSS laser (all from Coherent, Inc, San Jose, CA). Fluorescence was detected with a back-illuminated EMCCD camera (DU-897, Andor Technology) controlled by MetaMorph (Molecular Devices). Emission wavelengths were selected with filters (527/27 nm for GFP and 590 nm long-pass for mCherry) mounted in a filter wheel (Sutter Instruments). Islets and cells were mounted in an open perfusion chamber with temperature held at 37°C. For fluorescence recovery after photobleaching analysis of indicator mobility, a 3×3 μm area within the cells was exposed to a 100 ms 405-nm light pulse to bleach the fluorophores, followed by continued acquisition for 120 s at 4 fps. Calculations of the mobile fraction (*Mf*) of each fluorescent protein was performed using the following formula:

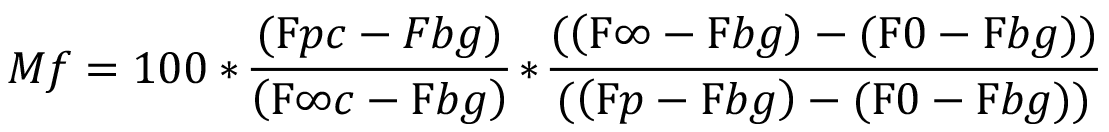

F*pc* (whole cell pre-bleach intensity), F*p* (bleach ROI pre-bleach intensity), F*∞c* (asymptote of fluorescence recovery of the whole cell), F*bg* (mean background intensity), F*∞* (asymptote of the bleach ROI), F*0* (bleach ROI post-bleach intensity).

#### Stimulated Emission Depletion (STED) microscopy

Sub-diffraction microscopy was performed with a Stedycon STED super-resolution instrument (Abberior Instruments, Germany) with an Olympus BX63F upright motorized microscope body. The objective was a 100x / 1.56 NA (Olympus, Japan) and the STED depletion laser had a wavelength of 775 nm. Emission filters were 575-625 nm and 650-700 nm corresponding to Star 580 and Star Red fluorophores, respectively.

### Mechanical stimulation of islets and cells

#### Islet compression

Islet compression experiments were performed with a custom-built manipulator referred to as the cell press. The device consists of 3D printed parts, a linear piezoelectric positioner (SLC-1730, SmarAct GmbH, Oldenburg, Germany), which is operated via a MCS2 manual controller (SmarAct), and a PDMS compression pillar, which was cast from a 3D-printed mold. The cell press was secured directly to the microscope behind the translation stage of an LSM700 confocal microscope (Zeiss, Jena, Germany). A relative zero position for the PDMS pillar was first established in an islet-free dish by bringing the surface of the pillar in contact with the bottom of the dish and recording the position displayed on the MCS2 controller. The pillar was retracted and a dish containing islets expressing Smo-GCaMP5G-mCherry were placed on the stage. Individual islets were imaged using a Plan-Apochromat 63x/1.4 (Zeiss) objective with the pinhole set to produce a 7.5 μm thick optical z-slice. Islets were imaged at their point-of-contact with the cover-glass and image acquisition was performed using Zen software (Zeiss). Islets were imaged prior to the initiation of compression experiments to establish a baseline for the GCamP5G and mCherry fluorescence signals. Compression experiments were performed by initially lowering the pillar to within approximately 200 – 300 μm from the zero position, after which the pillar was lowered in 50 μm steps until initial contact was made with the upper surface of the islet, after which compression continued in 5 μm steps. Compression continued until regions of the islet that were previously not visible (as they were above the imaged z-slice height) were pushed into the field of view, this point onward was referred to as deformation. This occasionally resulted in additional cilia being relocated into the field of view; however, image analysis was limited to cilia for which baseline GCamP5G and mCherry fluorescence signals had been established. Islets were compressed in the presence of 1.3 mM Ca^2+^ and where indicated were depolarized with 30 mM KCl.

#### Fluid flow stimulation

For mechanical stimulation of MIN6 cell cilia we used borosilicate pipets of 0.5-1μM diameter to puff extracellular solution locally (mM composition: NaCl 138, CaCl_2_ 2.6, MgCl_2_ 1.2, KCl 5.6, HEPES 10 and glucose 3; pH 7.4, 300 mOsm). The pipet solution was puffed toward the cilia by gently applying positive pressure with the mouth. A second borosilicated pipet with the same resistance was used to induce plasma membrane depolarization by puffing a solution containing (in mM): NaCl 63, CaCl_2_ 2.6, MgCl_2_ 1.2, KCl 8.6, HEPES 10 and glucose 3; pH 7.4, 300 mOsm. The mechanical stimulation pipet was located >5 μm from the cilia head and the high potassium pipet 10-15 μM from the cell body at the opposite side of the mechanical stimulation pipet. The mechanical stimulation was intermittently applied every 1s during 225s. During the recording, the cilia were allowed to rest from the stimulation 1 to 3 times to see possible changes at the cilia fluorescence baseline. Then, cells were let to rest 25s without stimulation before inducing membrane depolarization for 200s. The high potassium solution was puffed by applying constant positive pressure (3.7 psi) controlled by a Pico Spritzer (ValveLink8.2, Automate Scientific). The experiments were carried out at 32 °C. The samples were observed with a 40x Zeiss objective with a 1.2 NA. A Prime 95B CMOS camera (Teledyne Photometrics) was used to acquire the images, and GCaMP and mCherry fluorophores were excited by 473nm and 561 nm lasers (Cobolt, Stockholm, Sweden). Red and green channel images were acquired simultaneously at a 2Hz frequency, and the emission light was separated onto the two halves of the camera chip using an image splitter (Optical Insights) with a cutoff at 565nm (565dcxr, Chroma) and emission filters (FF01-523/610, Semrock; and ET525/50m and 600EFLP, both from Chroma). Cilia and cell bodies intensities over time were analyzed by homemade Macros in Fiji (ImageJ). Results are reported as the Δ(GCaMP/mCherry) intensity ratio.

### Immunofluorescence

Culture medium was removed and samples were washed with D-PBS at 37°C and fixed with 4% PFA in PBS for 5 min, then permeabilized with 0.5% TritonX-100 in PBS for 10 min and blocked with 2% BSA in PBS for 1 h. Primary antibodies were diluted in blocking solution and samples incubated for 2 h at RT (cells grown on coverslips were placed on a 100μl drop while 20~50 isolated islets were put into 200μl). Next, a wash with 2% BSA in PBS for 5 min was followed by 1 h incubation with secondary antibodies (same way as primary). Subsequently, samples were washed with 2% BSA in PBS for 5 min and mounted using ProLong™ Gold Antifade Mountant with DAPI (ThermoFisher Sci., USA). To preserve islets structural integrity, they were embedded into a 2% agar in PBS gel after being put into mounting medium.

### Image analysis

All images were analysed using the Fiji version of the ImageJ Software ^23^. Fluorescence line profiles were produced by drawing a segmented line over the cilium and part of cytoplasm for obtaining intensity values that were then scaled to the cytoplasmic or background levels (ratiometric data was obtained in a similar way by dividing profiles corresponding to different channels). Line width was set to match cilium diameter (approximately 3-5 pixels for confocal and TIRF, 7-9 pixels STED); for transverse profiles (fig 7H) line width was expanded to include the whole basal compartment (117 pixels). Analysis of live cell Ca^2+^ imaging data (TIRF) was generated by drawing ROIs containing the cilium from where mean values of pixel’s intensities were extracted from each time frame. Kymographs were created by using the corresponding function in Fiji; for ratiometric ones the GCaMP kymograph was divided by the corresponding mCherry kymograph. Center of mass calculation was performed on straightened cilia (Fiji function and analysis tool), the length coordinate was then normalized to total length of each individual cilium. The spontaneous Ca^2+^ activity in the cilium was analyzed using the *Neural Activity*^3^ (*NA^3^*) ^24^. This tool consists of a peak detection algorithm based in the continuous wavelet transform. The video is first divided in a grid which can be as dense as a single pixel per square, each zone in the grid is analyzed as an independent Ca^2+^ trace. Each trace is transformed using the Mexican-hat wavelet and the spectrum ridges are detected and grouped as a tree formation. Then those ridges are filtered in order to discard those of minor importance and keep the only the ones that contain possible activity peaks. Lastly the peaks in each of the remaining ridges are discarded based on the signal to noise ratio, leaving only those with a power over certain threshold. The phase-relationship between cilia and soma Ca^2+^ was determined using the *WaveletComp* package in R ^25^. A representative Ca^2+^ recording is selected for each structure, one for the cilia and one for the soma. Each of those traces is transformed with the continuous wavelet transform. This transformation results in a complex spectrum which contains information of magnitude and phase of the signals, which means that it is then possible to compare the phase spectrums of both signals and establish whether there exists a sustained phase relationship between both of them. That relationship is then presented in radians, see the package vignette for a thorough explanation.

### Statistical analysis

All data were processed using GraphPad Prism version 9.1.2 for Mac (GraphPad Software, San Diego, California USA, www.graphpad.com). Distribution of data sets was checked for normality and statistical test were chosen accordingly (see figure legends for details).

## RESULTS

### Measurements of cilia Ca^2+^ within intact pancreatic islets of Langerhans

Immunostaining of isolated mouse islets of Langerhans with an anti-acetylated tubulin antibody enabled visualization of primary cilia and revealed extensive ciliation. Most islet cells were equipped with a single cilium with a length of 5-10 μm (Fig. 1A and Suppl. Fig. 1). Co-staining for the two major hormones of the islets showed that both insulin-secreting β-cells and glucagon-secreting α-cells were ciliated (Fig. 1B, C). After 5 days of *in vitro* culture, most cells of the islet were positive for insulin and possessed prominent primary cilia, which were identified by immunostaining for IFT88 (Fig 1D, E), the well-characterized ciliary receptors Patched and Smoothened and the somatostatin receptor type 3 (SSTR3) (Fig. 1E). The enrichment of receptors on the cilia surface has led to the proposal that these structures function as antennae that sense the local environment, and Ca^2+^ has been proposed to be a relevant second messenger to propagate receptor signals within the cilium ^5,6^. We therefore developed a cilia-targeted Ca^2+^ biosensor, where Smoothened was used to confine the sensors expression to the cilium, and delivered it to islets through adenoviral transduction (Fig. 1F). More specifically, we fused the Ca^2+^ indicator GCaMP5G and the reference fluorophore mCherry to the C-terminus of Smoothened (Smo-GCaMP5G-mCh). The indicator was strongly enriched in the cilium, determined by co-staining for acetylated tubulin, and the topology of Smoothened in the cilia membrane was confirmed by acid quenching and proteinase K treatment, demonstrating that the fluorescent proteins localized to the cilia lumen (Suppl. Fig. 1).

**Figure 1.**
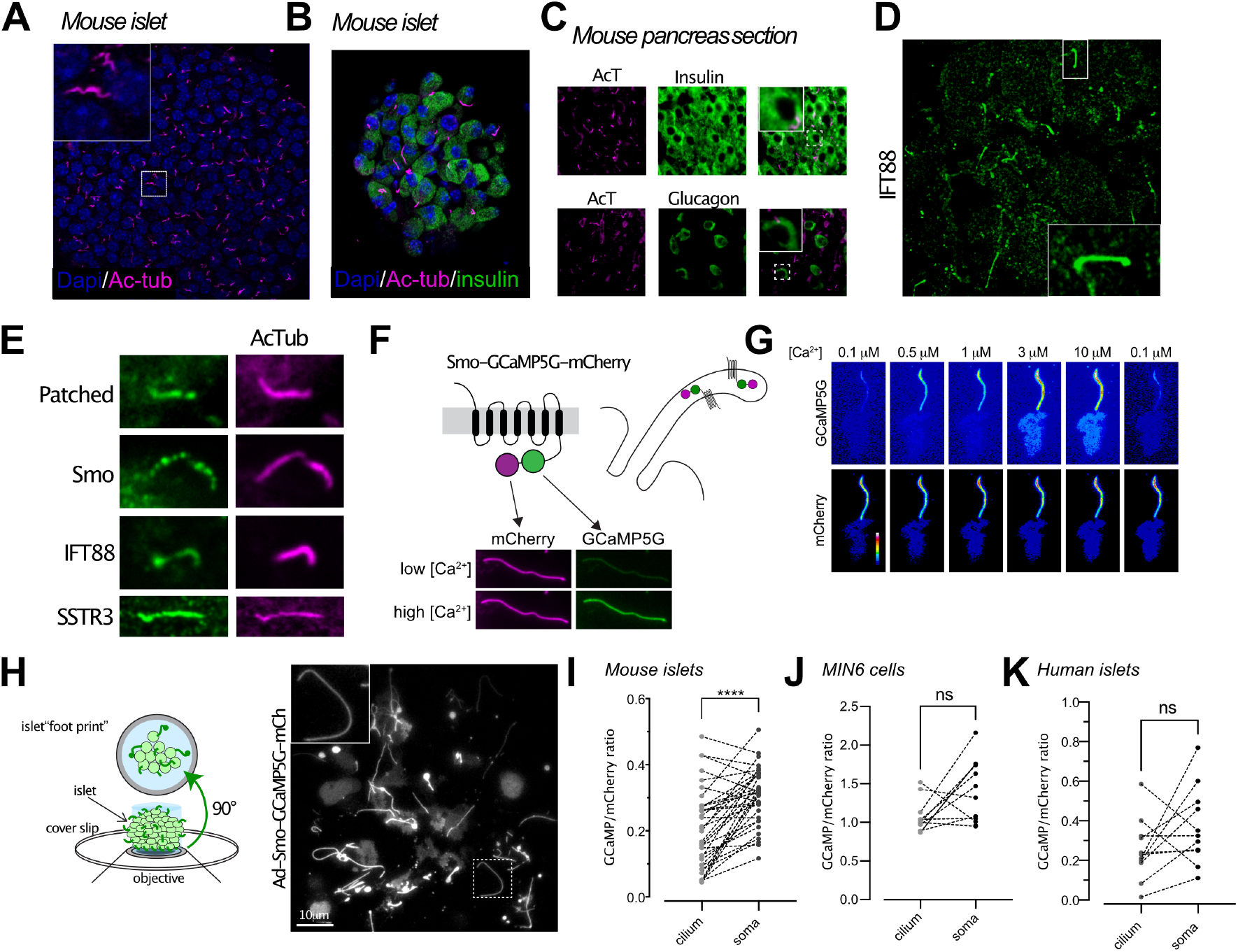
Ca^2+^ imaging in primary cilia of intact islets of Langerhans. A. Confocal microscopy image of a mouse islet where cilia are visualized by immunofluorescence staining against acetylated tubulin (magenta) and nuclei with Dapi staining (blue). B. Confocal microscopy image of a small mouse islet showing insulin in green, acetylated tubulin in magenta and nuclei in blue. C. Epifluorescence microscopy images of pancreas sections immunostained for acetylated tubulin (magenta), insulin (green, top panel) and glucagon (green, bottom panel). D. Confocal microscopy image of a mouse islet where cilia are visualized by immunofluorescence staining against IFT88 (green). E. Confocal micrographs of the localization of Patched, Smoothened, IFT88 and SSTR3 (all green) to the primary cilium (magenta) of mouse islet cells. F. Schematic illustration showing the principle of the ratiometric cilia-targeted Ca^2+^ indicator Smo-GCaMP5G-mCh. G. TIRF microscopy images of Smo-GCaMP5G-mCherry fluorescence from MIN6 cells. The cells were permeabilized with α-toxin and exposed to the indicated Ca^2+^-buffer. Top row shows fluorescence change of GCaMP5G in response to the different Ca^2+^ buffers, while the bottom row shows the corresponding change in mCherry fluorescence. H. Principle of TIRF microscopy imaging of primary cilia in intact islets of Langerhans. I. Quantifications of the resting GCaMP5G/mCherry fluorescence in the cilia and soma of mouse islet cells shows that the Ca^2+^ concentration is lower in the cilium (n=39; *** P<0.001; paired, 2-tailed Student’s t-test). J. Quantifications of the resting GCaMP5G/mCherry fluorescence in the cilia and soma of MIN6 cells shows that there is no difference in Ca^2+^ concentration between the two compartments (n=12; 2-tailed Student’s t-test). K. Quantifications of the resting GCaMP5G/mCherry fluorescence in the cilia and soma of human islet cells shows that there is no difference in Ca^2+^ concentration between the two compartments (n=11; paired, 2-tailed Student’s t-test).

To evaluate the properties of this Ca^2+^-indicator, we permeabilized clonal MIN6 β-cells expressing Smo-GCaMP5G-mCh and perifused the cells with Ca^2+^-buffers while recording GCaMP5G and mCherry fluorescence using TIRF microscopy. We observed stepwise increases in ciliary Ca^2+^, with half-maximal and maximal effect at 0.6 μM and 3 μM, respectively (Fig. 1G and Suppl. Fig. 1). Although the majority of Smo-GCaMP5G-mCh was localized to the cilium, small amounts of the protein were also found in the plasma membrane, which enabled direct comparisons of GCaMP5G fluorescence change in the two compartments. Importantly, the indicator localization did not affect the response to Ca^2+^, and the observed EC_50_ was also similar to that reported previously for soluble GCaMP5G ^21^ (Suppl. Fig. 1). The expression of Smo-GCaMP5G-mCh resulted in cilia lengthening and the appearance of bulky distal tips (Fig. 1H and Suppl. Fig. 1), likely indicating exacerbated Hedgehog signaling when over-expressing Smoothened. Photolysis of caged Ca^2+^ in the cytosol of individual MIN6 cells expressing Smo-GCaMP5G-mCh resulted in a Ca^2+^ wave that propagated into the cilium, and we did not notice any correlation between indicator expression level and the kinetics of the Ca^2+^ response in the cilium, suggesting that the expression of the indicator has negligible buffering effect (Suppl. Fig. 1). We also performed photobleaching experiments to determine the mobility of the indicator in the ciliary and plasma membranes, and found that the indicator was essentially immobile in both compartments (Suppl. Fig. 1).

Ca^2+^ is considered an important second messenger in the cilium despite the fact that its actual role is still debated and poorly understood ^26,27^. One key aspect is how the resting concentration in the cilium compares to that of the cytoplasm, and estimations based on electrophysiological and imaging approaches indicate that the ciliary Ca^2+^ concentration is higher than the cytosolic Ca^2+^ concentration ^5^. We evaluated the relative concentrations in the two cellular compartments by comparing the average ratios of fluorescence line profiles drawn over the cilium and part of plasma membrane, as visualized by TIRF microscopy, in both mouse and human islet β-cells and in clonal MIN6 β-cells expressing Smo-GCaMP5G-mCh (Fig. 1I-K). In contrast to previous studies, we found that the GCaMP5G-to-mCherry ratios in the cilia were either the same, or slightly lower, than those of the cytosol, indicating a lower resting Ca^2+^ concentration in the cilia. Control experiments showed that this is likely not due to acid quenching of the pH-sensitive GCaMP5G, since a cilia-targeted pH indicator revealed that the cilia lumen is slightly more alkaline than the cytosol (Suppl. Fig. 1).

### The primary cilium is isolated against cytosolic Ca^2+^ concentration changes

Next, we determined what effect changes in the cytosolic Ca^2+^ concentration has on Ca^2+^ concentrations in the primary cilium of cells within intact islets. To this end, cytosolic Ca^2+^ was increased in three different ways; by depolarization-induced entry from the extracellular space, by release from intracellular stores, and by photorelease of caged Ca^2+^, while the Ca^2+^ concentration changes were recorded using Smo-GCaMP5G-mCh and TIRF microscopy. This technique enabled simple access to primary cilia protruding from the apical surfaces of the outer cell layer of the islet, and also enabled extended recordings of cilia Ca^2+^ thanks to low photobleaching and immobilization of cilia beneath the weight of the islet. The depolarization-induced opening of voltage-gated Ca^2+^ channels in islet β-cells caused a robust increase in the cytosolic Ca^2+^ concentration (Fig. 2A-C). We segmented the cilium into sub-compartments and observed a small rise of Ca^2+^ at the base of the cilium which mirrored the change in cytosolic Ca^2+^, while more distal compartments of the cilium instead exhibited a drop in GCaMP5G fluorescence, indicative of lowering of the Ca^2+^ concentration (Fig. 2B). This indicates the existence of a Ca^2+^ diffusion barrier and mechanisms operating at the base of cilium that prevents the penetration of Ca^2+^ from the cytosol into the cilium. An alternative source of Ca^2+^ is represented by internal stores. Release of Ca^2+^ from the endoplasmic reticulum (ER) was triggered by the muscarinic receptor agonist carbachol while islets were kept in 20 mM glucose to keep the ER-stores filled and diazoxide and verapamil to prevent glucose-induced Ca^2+^ influx. As expected, activation of muscarinic receptors produced a fast transient increase in cytosolic Ca^2+^ (Fig 2D), which was accompanied by an even more pronounced ciliary Ca^2+^ increase that, however, was restricted to the base of the cilium (Fig. 2E, F).

**Figure 2.**
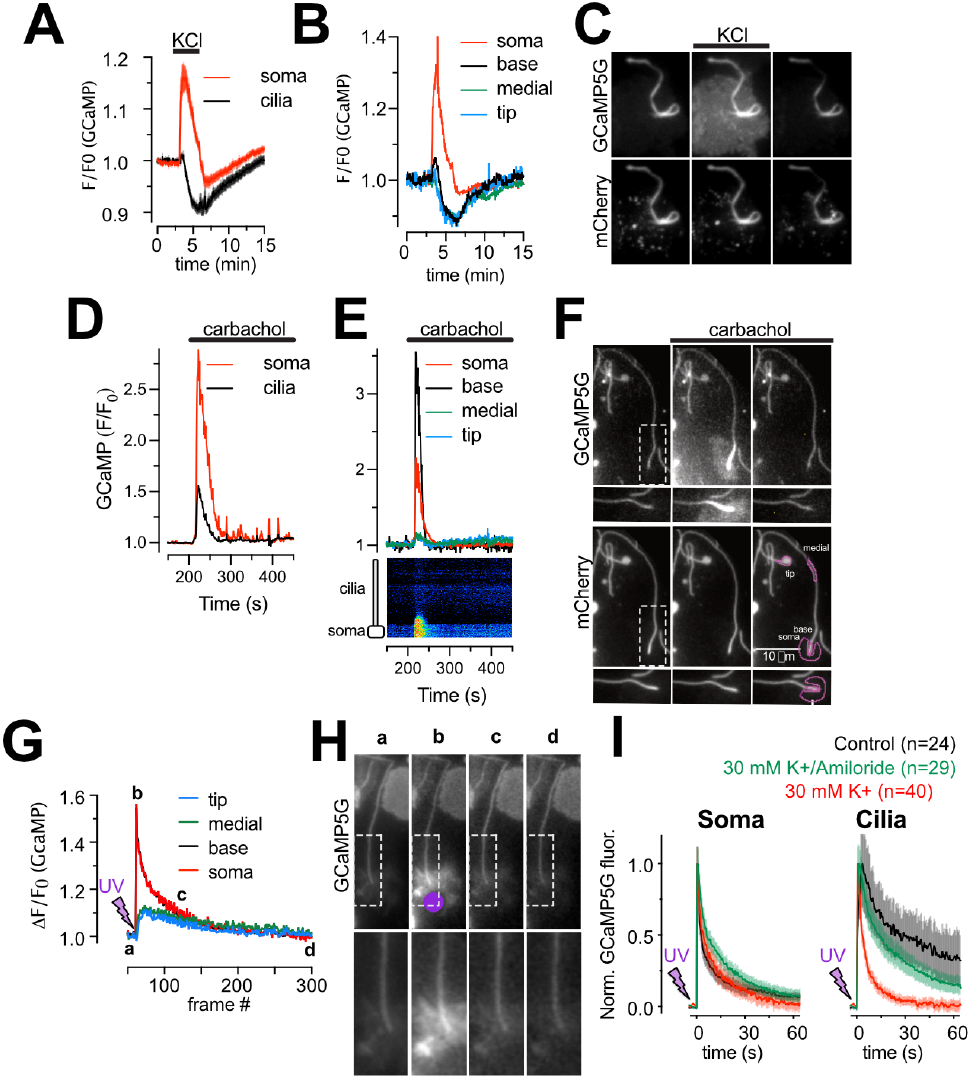
Restricted entry of cytosolic Ca^2+^ into the primary cilium of islet cells. A. TIRF microscopy recordings of GCaMP5G fluorescence from mouse islet cells transduced with an adenovirus encoding Smo-GCaMP5G-mCh. The islet was exposed to a brief depolarization (30 mM KCl) while the Ca^2+^ concentration was recorded in the soma and cilium. Data presented are averages (±S.E.M.) from 10 cells from 1 islet and representative of 15 islets. B. Ca^2+^ concentration change along the length of a mouse islet cell cilium during depolarization with 30 mM KCl, shown as the relative GCaMP5G fluorescence change at three distinct segments (base, medial and tip). C. TIRF microscopy images of Smo-GCaMP5G-mCh fluorescence change during KCl-depolarization. Notice the lack of Ca^2+^ increase in the more distal parts of the cilium. D. TIRF microscopy recordings of GCaMP5G fluorescence from mouse islet cells transduced with an adenovirus encoding Smo-GCaMP5G-mCh. The islet was exposed to a 10 μM carbachol while the Ca^2+^ concentration was recorded in the soma and cilium. Data presented are averages (±S.E.M.) from 12 cells from 1 islet and representative of 15 islets. E. Ca^2+^ concentration change along the length of a mouse islet cell cilium during carbachol stimulation, shown as the relative GCaMP5G fluorescence change at three distinct segments (base, medial and tip). Kymograph from the same cell is shown below. F. TIRF microscopy images of Smo-GCaMP5G-mCh fluorescence change during carbachol stimulation. Notice the prominent rise of Ca^2+^ at the base of the cilium, but the lack of spreading to more distal segments. G. Ca^2+^ concentration change along the length of a mouse islet cell cilium in response to photolysis of caged-Ca^2+^, shown as the relative GCaMP5G fluorescence change at three distinct segments (base, medial and tip). H. TIRF microscopy images of Smo-GCaMP5G-mCh fluorescence change following Ca^2+^-uncageing (purple dot indicate site of photolysis). Notice the prominent rise of Ca^2+^ at the base of the cilium, but the lack of spreading to more distal segments. I. Recordings of GCaMP5G fluorescence change in the soma and cilia of mouse islet cells under control conditions (black) and when islets were exposed to 30 mM KCl (red) or 30 mM KCl in combination with Amiloride (green). Data presented as means ± S.E.M. for the indicated number of cells.

Next, we triggered acute elevations of cytosolic Ca^2+^ by photorelease of caged Ca^2+^ in discrete cytosolic regions at distance from the cilium (Fig 2G, H). The photorelease generated a Ca^2+^ wave that propagated into the cilium. Interestingly, the spreading was strongly dampened within the cilium and was further reduced under conditions when the islets were depolarized by 30 mM KCl. This effect of depolarization was lost when 200 μM amiloride was included in the solution. Importantly, neither depolarization nor amiloride affected the kinetics of the Ca^2+^ response in the cytosol (Fig. 2I). Depolarization therefore appears to enhance Ca^2+^ extrusion from the cilium while amiloride abolished the enhancement, likely by blockade of the Na^+^/Ca^2+^-exchanger.

The primary cilium is a mechanosensory organelle in many cell types, although to what extent Ca^2+^ is a relevant signaling molecule downstream of mechanical stimulation in cilia is controversial ^26,27^. We therefore exposed islet cells to two forms of mechanical stimulation, mild compression and fluid flow, while recording Ca^2+^ concentration changes in the primary cilia by confocal microscopy. Compression was accomplished using the cell press, a piezomotor-controlled manipulator that linearly translates a flexible PDMS compression pillar, which was fitted to the stage of the microscope. This device enabled fine-tuned control of intra-islet pressure (Fig. 3A, B), and neither weak (which produced no morphological change in islet structure) nor strong (which caused visible islet deformation) compression resulted in dramatic increases in cilia Ca^2+^ concentration (Fig. 3B). However, mild compression did elicit a rise of Ca^2+^ in the cilia of some islet cells (16/161 cilia, P<0.0001) (Fig. 3C, D). Since compression affects both cilia and soma of the islet cells, it is possible that the observed effect is secondary to the compression of the soma. To directly stimulate cilia without affecting the cell bodies, we exposed MIN6 β-cell cilia to fluid flow delivered through a pulled glass pipette placed next to a cilium using a micromanipulator (Fig. 3E). Local “puffing” of the bath solution onto individual cilia caused noticeable movement of the structure but did not result in any change in cilia Ca^2+^ concentration (Fig. 3F, G). Together, these results suggest that β-cell cilia do not respond directly to mechanical stimulations with increased levels of Ca^2+^.

**Figure 3.**
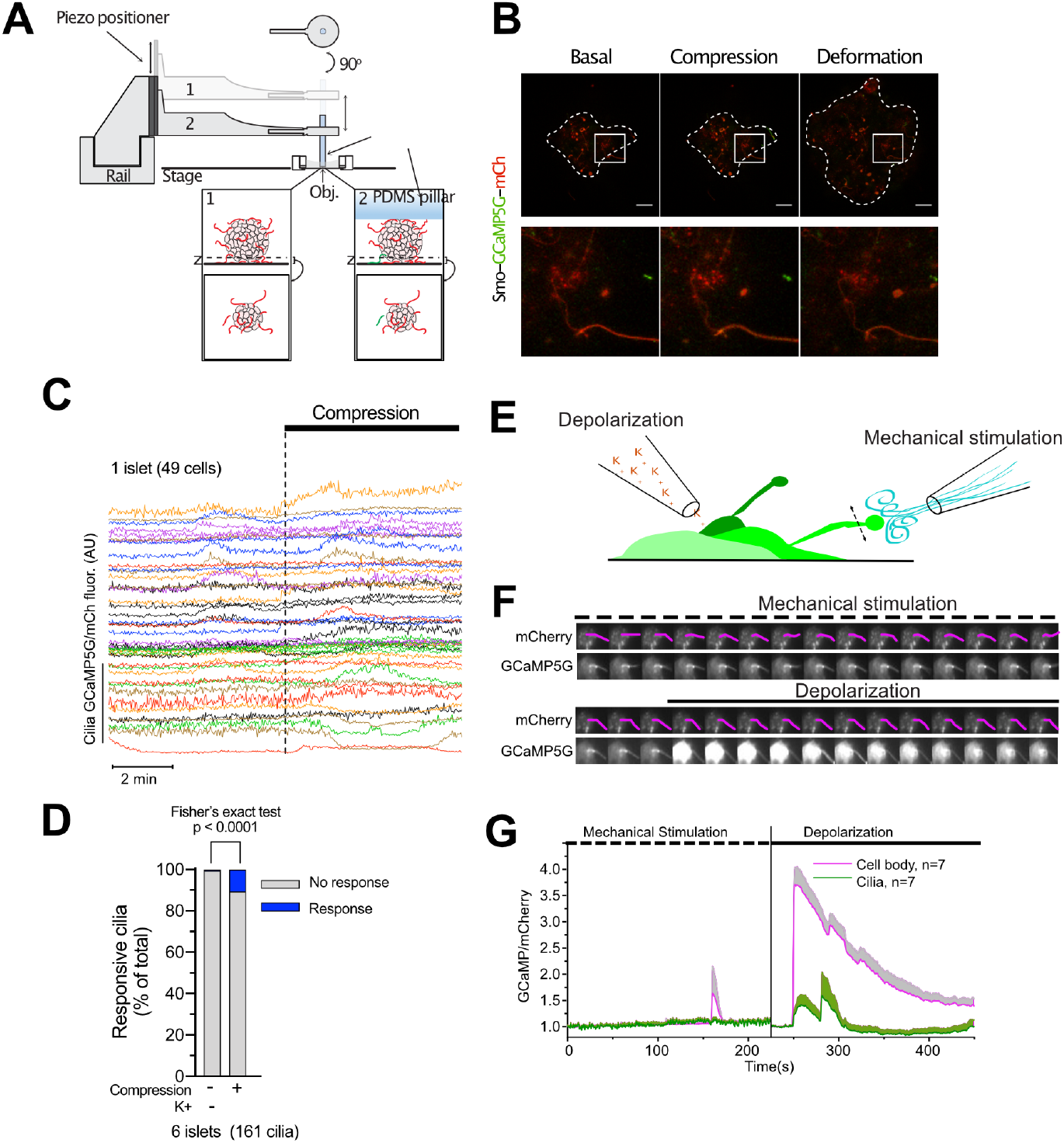
Cilia Ca^2+^ measurements in mechanically stimulated islet cells. A. Schematic drawing of the cell-press used to compress islets on the stage of a confocal microscope. B. Confocal micrographs of a whole islet (top) and a magnified section of the same islet (inset below) expressing Smo-GCaMP5G-mCh. Notice how both mild compression and deformation is without effect on both cilia and cytosolic Ca^2+^. C. Changes in cilia GCaMP5G/mCh fluorescence ratio during mild compression of a mouse islet (n=49 cells). D. Fraction of islet cell cilia exhibiting Ca^2+^ increases after mild islet compression (n = 161 cilia from 6 islets). F. Confocal microscopy images of a MIN6 cell expressing Smo-GCaMP5G-mCh following exposure to mechanical stimulation (top) and depolarization (bottom). G. Means ± S.E.M. (n=7) for the cilia and cell body GCaMP5G fluorescence change in response to mechanical stimulation and depolarization. The increase in cilia fluorescence upon depolarization is likely a contamination from the cytoplasm, which partially overlap with the cilia in these recordings.

### Cilia Ca^2+^ dynamics in glucose-stimulated islet cells

To determine the cilia Ca^2+^ activity in a physiological context, we exposed Smo-GCaMP5G-mCh-expressing islets to a buffer containing 11 mM glucose, a concentration at which insulin secretion is strongly stimulated. This resulted in an immediate and very pronounced increase in GCaMP5G fluorescence in the islet cell cilia (Fig. 4A and Suppl. Fig. 2). Less pronounced increases were also seen in the cell bodies. This rise occurred before depolarization-induced Ca^2+^ influx and was not seen with soluble GCaMP ^28^ (Suppl. Fig. 2), making us question whether it reflects a Ca^2+^ concentration change. One possibility is that ATP generated by glucose metabolism affects the fluorescent properties of GCaMP5G ^29^. Experiments performed in permeabilized MIN6 cells expressing Smo-GCaMP5G-mCh or a plasma membrane-anchored red Ca^2+^ indicator (Lyn_11_-R-GECO) did however not reveal any direct impact of ATP on either GCaMP5G or R-GECO fluorescence (Suppl. Fig. 2). Measurements of pH using 5HT_6_-venus-CFP ^6^ showed that glucose-stimulation alkalized both cytosol and cilia. Since GCaMP5G fluorescence is strongly influenced by pH ^30^, it is reasonable to assume that the initial glucose-induced increase in GCaMP5G fluorescence is caused by a rise in pH (Suppl. Fig. 2). This glucose-induced rise of GCaMP5G fluorescence was followed by the appearance of cytosolic Ca^2+^ oscillations with a period of 2-5 min (Fig. 4A). We observed similar periodic fluctuations in the cilia Ca^2+^ concentration, and when we analyzed the low frequency component of the oscillations in pairs of connected cilia and cell bodies, we found a stable phase shift of -π/2 between the two signals (Fig. 4A-B). The anti-phase locking of the cilia and cytosolic Ca^2+^ oscillations may reflect the enhancement of Ca^2+^ extrusion due to depolarization taking place at supra-threshold membrane potentials that marks the peak of cytoplasmic [Ca^2+^] and the nadir of the ciliary [Ca^2+^]. Inspection of the change in cilia Ca^2+^ concentration along the length of the cilium revealed a pattern reminiscent of that observed following depolarization, where the cytosol-proximal region exhibited changes similar to the cytosol (i.e., oscillations in-phase), whereas more distal regions were phase-shifted (Fig. 4C). There was a positive correlation between the duration of the cytosolic [Ca^2+^] increase and the decrease in cilia [Ca^2+^] (Fig. 4D, E), and pulsatile application of depolarizing KCl concentrations mimicked the effect of glucose (Fig. 4F). Addition of the K_ATP_-channel opener diazoxide or the L-type voltage-dependent Ca^2+^ channel inhibitor methoxyverapamil inhibited the glucose-induced Ca^2+^ oscillations in both compartments (Fig. 4G, H), while the addition of the cAMP-elevating agent forskolin transformed the slow cytosolic Ca^2+^ oscillations to rapid spike-like fluctuations and was accompanied by the disappearance of cilia Ca^2+^ oscillations (Fig. 4I). Together, these results indicate that the glucose-induced Ca^2+^ concentration changes in the cilium and soma are driven by a common underlying mechanism. The cilium is devoid of mitochondria but has been shown to have intrinsic glycolysis and could therefore potentially generate local ATP from glucose that might drive Ca^2+^ influx or extrusion ^31^. To test if a cilia-derived messenger might be responsible for the observed anti-synchronicity, we exposed islets to the mitochondrial substrate α-ketoisocaproic acid (5 mM). This resulted in similar slow Ca^2+^ oscillations in the β-cell cilia that were out-of-phase with those of the cell bodies (Fig. 4J). Taken together, these results confirm our previous observations that the islet cell cilia are effectively isolated against changes in the cytosolic Ca^2+^ concentration.

**Figure 4.**
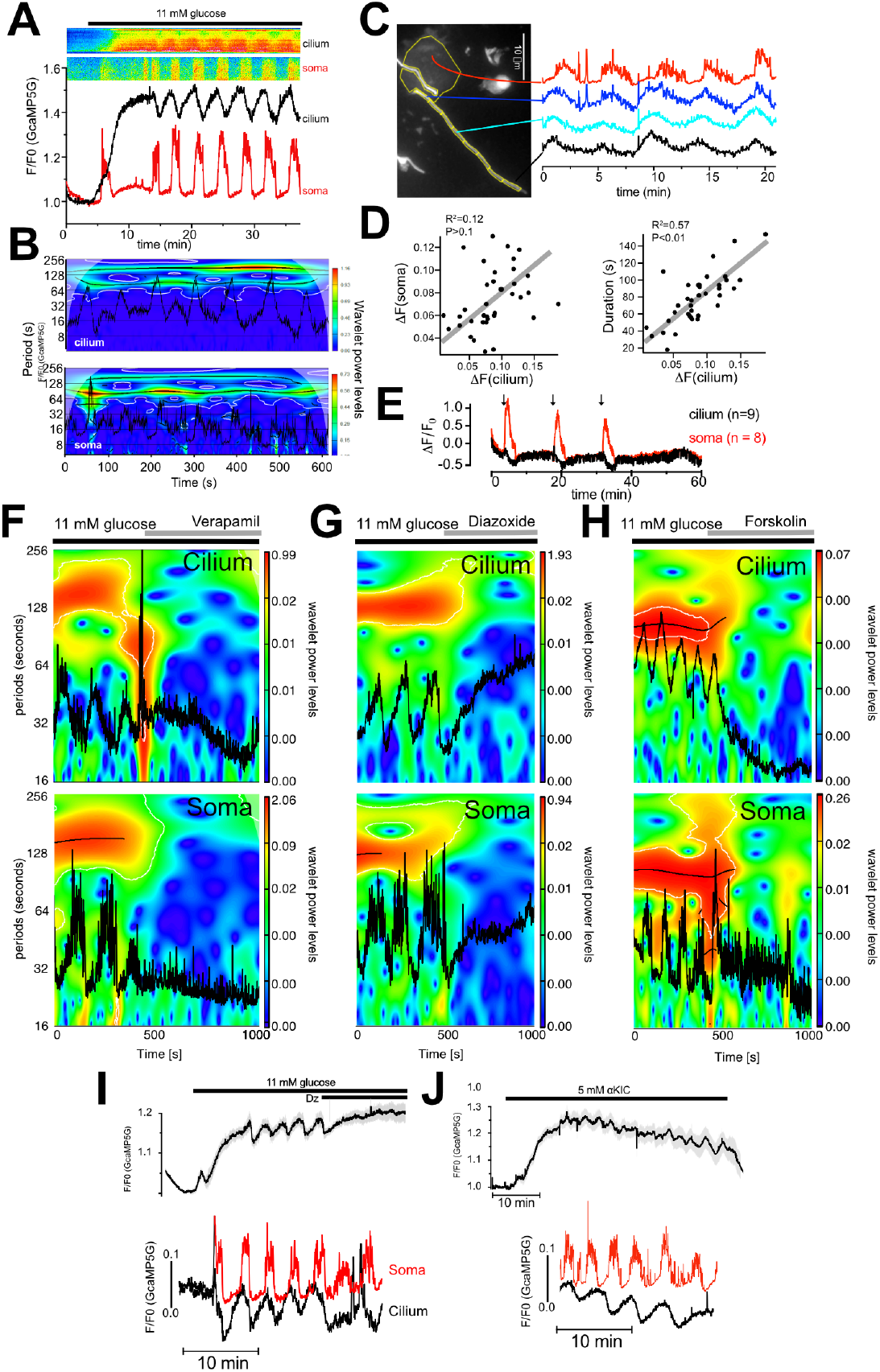
Cilia and cytosolic Ca^2+^ are regulated by a common mechanism. A. TIRF microscopy recording of Smo-GCaMP5G-mCh fluorescence from a β-cell within an intact mouse islet during an increase in the glucose concentration from 3 mM to 11 mM. Notice the appearance of regular oscillations in both cilium and soma. Kymograph above shows GCaMP5G/mCh ratio changes in a small part of the soma and along the cilium. B. Color coded spectrogram of ciliary (top) and somatic (bottom) frequency components with superimposed traces (black) of the whole compartments Ca^2+^ activity recorded from one cell during 11 mM glucose exposure. Note the prevalence of a slow component with a period of about 100 s in both cilium and soma. C. GCaMP5G fluorescence change from a mouse islet cell stimulated with 11 mM glucose. The cilium has been segmented and the fluorescence change within the segments are shown to the right. Notice that there is a gradual change of the oscillatory pattern as the segments becomes more distal to the cell body. D. Scatter plots showing a positive correlation between the relative Ca^2+^ concentration changes in the cilia and soma (left) and between the duration of the cytosolic Ca^2+^ increase and the change in cilia Ca^2+^ concentration (right). E. TIRF microscopy recording of GCaMP5G fluorescence change in the cilium (black) and soma (red) of islets cells following exposure to three 3 min KCl-depolarizations (arrows). Data are means ± S.E.M. for 8 cells from 1 islet and representative of 15 islets. F-H. Color coded power spectra of ciliary (top) and somatic (bottom) frequency components with superimposed traces (black) of the whole compartments Ca^2+^ activity recorded from individual cells exposed to different treatments (see horizontal lines above plots). I. TIRF microscopy recording of GCaMP5G fluorescence from 11 cilia from 1 mouse islet (means ± S.E.M.) during elevation of the glucose concentration from 3 to 11 mM and subsequent addition of 250 μM diazoxide (Dz). Shown below are recordings of cilia and cytosolic Ca^2+^ from a matched cilia-soma pair within the islet. Representative of 7 islets. J. TIRF microscopy recording of GCaMP5G fluorescence from 12 cilia from 1 mouse islet (means ± S.E.M.) following addition of 5 mM α-KIC. Shown below are recordings of cilia and cytosolic Ca^2+^ from a matched cilia-soma pair within the islet. Representative of 8 islets.

### Spontaneous cilia Ca^2+^ flashes in isolated pancreatic islets

The evidence presented so far suggests that the cilium is effectively isolated and actively purged of any interfering cytoplasmic Ca^2+^ attempting to enter the organelle. Such a protected status could be relevant in the event of locally generated ciliary Ca^2+^ signals. Prolonged recordings of cilia Ca^2+^ concentrations performed in mouse islets kept in a low (3 mM) glucose-containing buffer revealed the occurrence of spontaneous ciliary Ca^2+^ flashes (Fig. 5A). Similar flashes were also observed in clonal MIN6 β-cells and in human islet cells (Fig. 5B, C and Suppl. Fig. 3). Flashes were characterized by fast onset and offset transitions and it was possible to identify the site of origin and to follow Ca^2+^ spreading along the cilium (Fig. 5D-I). Occasionally it was even possible to detect a small rise of cytosolic Ca^2+^ as the cilia Ca^2+^ wave reached the base (see Fig. 5A). The propagation of Ca^2+^ in the cilium had a similar diffusion rate as that previously reported ^5^, and the average duration of these events was close to 1 minute (Fig. 5E-G). Notably, most of the flashes started in the more distal regions of the cilium, including the tip. Observations of cilia Ca^2+^ flashes throughout the cilia of an islet revealed a degree of temporal coordination, but not perfect synchronization, between the events across the islet, which may indicate that these fluctuations are caused by the action of a soluble molecule rather than electrical coupling (Fig. 5J). Flashes were suppressed when the glucose concentration was elevated or when islet cells were depolarized with KCl (Fig. 5K-M). Interestingly, the suppression of flashes induced by high glucose was abolished when the hyperpolarizing agent diazoxide and the voltage-dependent Ca^2+^ channel inhibitor verapamil were added (Suppl. Fig. 3), suggesting that cilia Ca^2+^ entry is not mediated by voltage sensitive channels and that it is suppressed at more positive membrane potentials. Blocking the Na^+^/Ca^2+^-exchanger with amiloride increased the duration of the events, suggesting that extrusion is involved in shaping the flashes (Suppl. Fig. 3).

**Figure 5.**
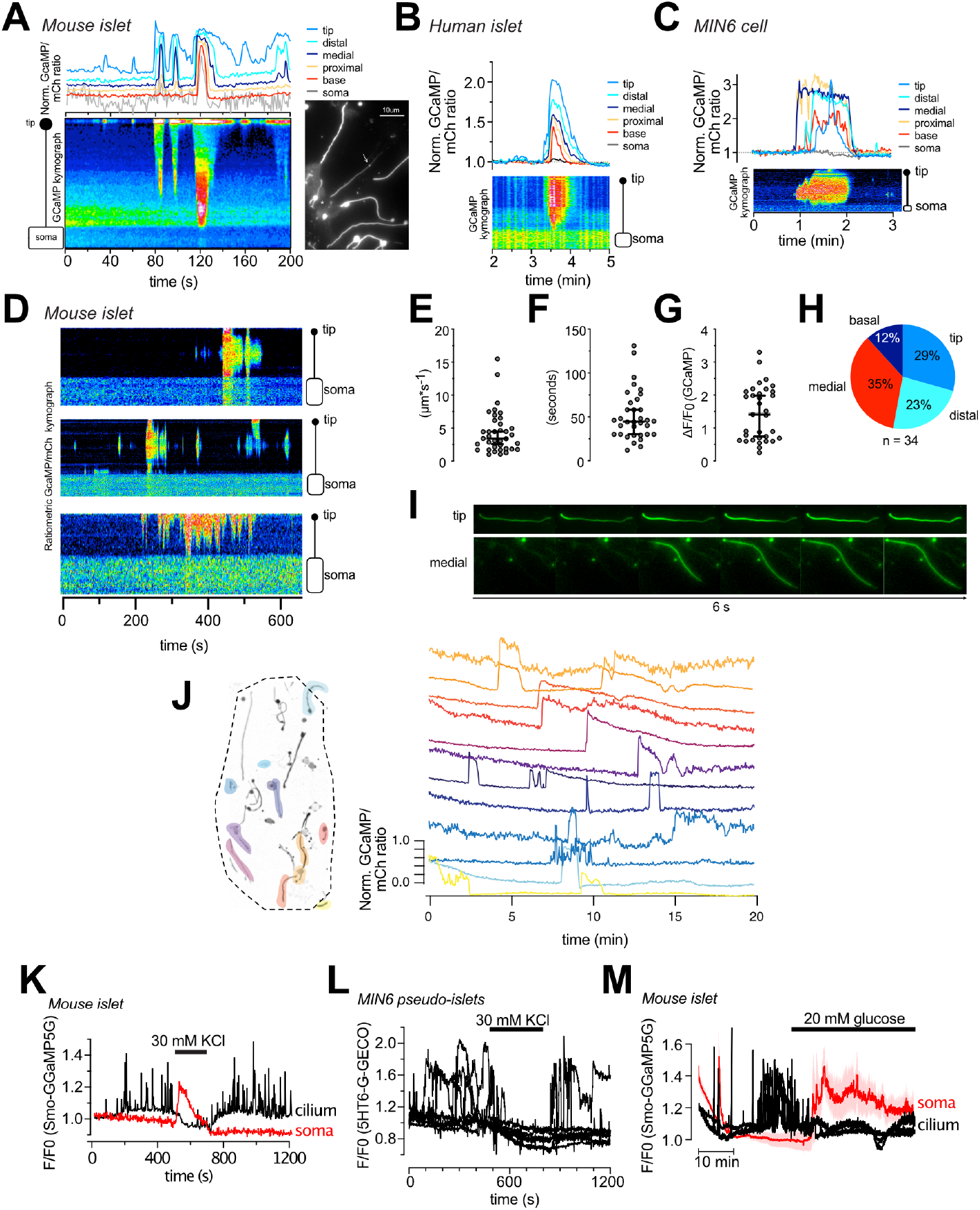
Isolated cilia Ca^2+^ activity within intact islets of Langerhans. A. TIRF microscopy recording of CaMP5G fluorescence from the soma and cilium of a mouse islet cell during a spontaneous Ca^2+^ “flash” (white arrow points to the cilium). Traces show the Ca^2+^ concentration change at different segments of the cilium and in the soma (grey) and the kymograph below shows the GCaMP5G fluorescence change along a line going from the soma to the tip of the cilium. B. A spontaneous cilia Ca^2+^ flash recorded from a human islet β-cell expressing Smo-GCaMP5G-mCh. C. A spontaneous cilia Ca^2+^ flash recorded from a MIN6 β-cell expressing Smo-GCaMP5G-mCh. D. Kymographs showing examples of spontaneous Ca^2+^ activity in three cilia-soma pairs within intact mouse islets. E. Ca^2+^ diffusion rates in the primary cilium of mouse islet cells during spontaneous Ca^2+^ flashes (n=35 cilia from 30 islets). F. Duration of spontaneous Ca^2+^ flashes in mouse islet primary cilia (n=35 cilia from 30 islets). G. GCaMP5G fluorescence change during spontaneous Ca^2+^ flashes in mouse islet cells (n=35 cilia from 30 islets). H. Site of Ca^2+^ flash origin in the primary cilium (n=34 cilia). I. Examples showing the initiation and propagation of Ca^2+^ along the cilium of islet cells. J. GCaMP5G fluorescence change in the primary cilia of a mouse islet kept in a basal buffer containing 3 mM glucose. Picture to the left shows the distribution of cilia within the islet (color represents individual cilia. Unmarked cilia did not exhibit spontaneous activity during the recording time). Traces to the right show the fluorescence change in individual cilia over time. Notice that there is certain coordination in responses between cilia of the islet. K. TIRF microscopy recording of GCaMP5G fluorescence in the cilium and cell body of an islet cell exposed to brief depolarization by 30 mM KCl. Notice that the rise of cytosolic Ca^2+^ is accompanied by the suppression of cilia Ca^2+^ flashes. L. TIRF microscopy recording of GCaMP5G fluorescence in the cilia of MIN6 cells aggregated into pseudo-islets and exposed to brief depolarization by 30 mM KCl. Notice that the rise of cytosolic Ca^2+^ is accompanied by the suppression of cilia Ca^2+^ flashes. M. TIRF microscopy recording of GCaMP5G fluorescence in the cilia and cell body of islet cells exposed to 20 mM glucose. Notice that the rise of cytosolic Ca^2+^ is accompanied by the suppression of cilia Ca^2+^ flashes (means ± S.E.M. for 10 cell bodies; 3 individual cilia within the same islet).

### GABA initiate localized cilia Ca^2+^ signalling via GABA-B receptors

To identify molecules that could initiate ciliary Ca^2+^ activity we exposed MIN6 cells expressing Smo-GCaMP5G-mCh to GPCR agonists known to be present within islets. We found that the addition of GABA at low concentrations (1-10 nM) caused activation of ciliary Ca^2+^ signaling but was without effect on the cytosolic Ca^2+^ concentration (Fig 6A-C). Baclofen, a selective agonist for metabotropic GABA-B receptors, had a similar effect as GABA, indicating that the evoked responses were triggered by activation of GABA-B receptors (Fig. 6C). We next tested the effect of GABAergic activation on cilia Ca^2+^ signaling in mouse islets. To quantify ciliary Ca^2+^ activity, we counted the number of events (defined as an increase in GCaMP5G fluorescence above 25% of resting levels) and calculated their density over defined periods of time (Fig. 6D). Baclofen increased ciliary Ca^2+^ activity and vigabatrin, which blocks GABA catabolism and promotes endogenous GABA accumulation and release, had a similar effect (Fig. 6E, F and I). Antagonists for both ionotropic (picrotoxin) and metabotropic (CGP 35348) GABA receptors did not suppress spontaneous activity, but their washout led to a significant increase of events (Fig. 6G, I). We also tested the effect of prolonged incubation (>5 hours) with vigabatrin and observed a trend towards suppression of cilia Ca^2+^ flashes, which might indicate exhaustion of GABA release or receptor desensitization (Fig. 6H, I). Having shown that activation of GABA receptors triggers Ca^2+^ activity in the cilium of β-cells, we determined the distribution of GABA receptors in islet cells by immunostaining. We first validated the selectivity of the GABA-B1 receptor antibody using siRNA knock-down (KD) in MIN6 cells (Suppl. Fig. 4). Control cells showed the presence of the receptors throughout the cilium while in KD cells the signal was significantly reduced. The localization pattern of GABA-B1 receptor in isolated mouse islets of Langerhans was strikingly different from MIN6 cells and the receptor was instead restricted to a short segment of the proximal cilium (Fig. 7A, B and D). Similar results were obtained using an antibody targeting an intracellular epitope of the GABA-B1 receptor (Fig. 7B). We did not detect any enrichment of GABA-A receptors or GABA-B2 receptors in the primary cilium (Fig. 7C, D). High resolution imaging using a STED microscope confirmed the confinement of GABA-B1 receptors at the base of the cilium (Fig. 7E). We characterized the receptor distribution by calculating the center of mass of straightened images and as expected the acetylated tubulin center of mass was close to half the length of the cilium while that of GABA-B1 was skewed towards the base (Fig. 7F, G). Transverse profiles showed a hollow signature of GABA-B1 which is considered to be a hallmark of periaxonemal particles, including receptors at the ciliary membrane (Fig. 7H). Importantly, expression of Smo-GCaMP5G-mCh did not alter GABA-B1 receptor distribution in the cilium (Fig. 7D, E). Receptor accumulation at the base of the cilium is compatible with two models; either these are receptors exiting the cilium following activation or they are receptors trapped in this compartment, ready to be mobilized into more distal segments of the organelle. Two facts support the latter model; first, most of the spontaneous Ca^2+^ activity and GABA-induced activity originated in the distal parts of the cilium (see Fig. 6), and second the inversin compartment has been hypothesized to work by sequestering ciliary proteins and releasing them under appropriate stimulation ^32^. To test this, we analyzed the GABA-B1 receptor distribution after a short (10 min) stimulation with GABA or the GABA-B receptor agonist baclofen. We found that both GABA and Baclofen caused a lengthening of the GABA-B1 positive segment at the base of cilia in the mouse islet (Suppl. Fig. 4). Consistent with this observation, the center of mass of GABA-B1 receptor distribution in STED images shifted towards the distal ciliary region and we could observe clusters of GABA-B1 receptors along the whole cilium (Fig. 7I). These observations strengthen the idea that GABA-B1 receptors are confined to a compartment at the base of the cilium and that signaling putatively involves diffusion of the receptor into more distal segments of the cilium.

**Figure 6.**
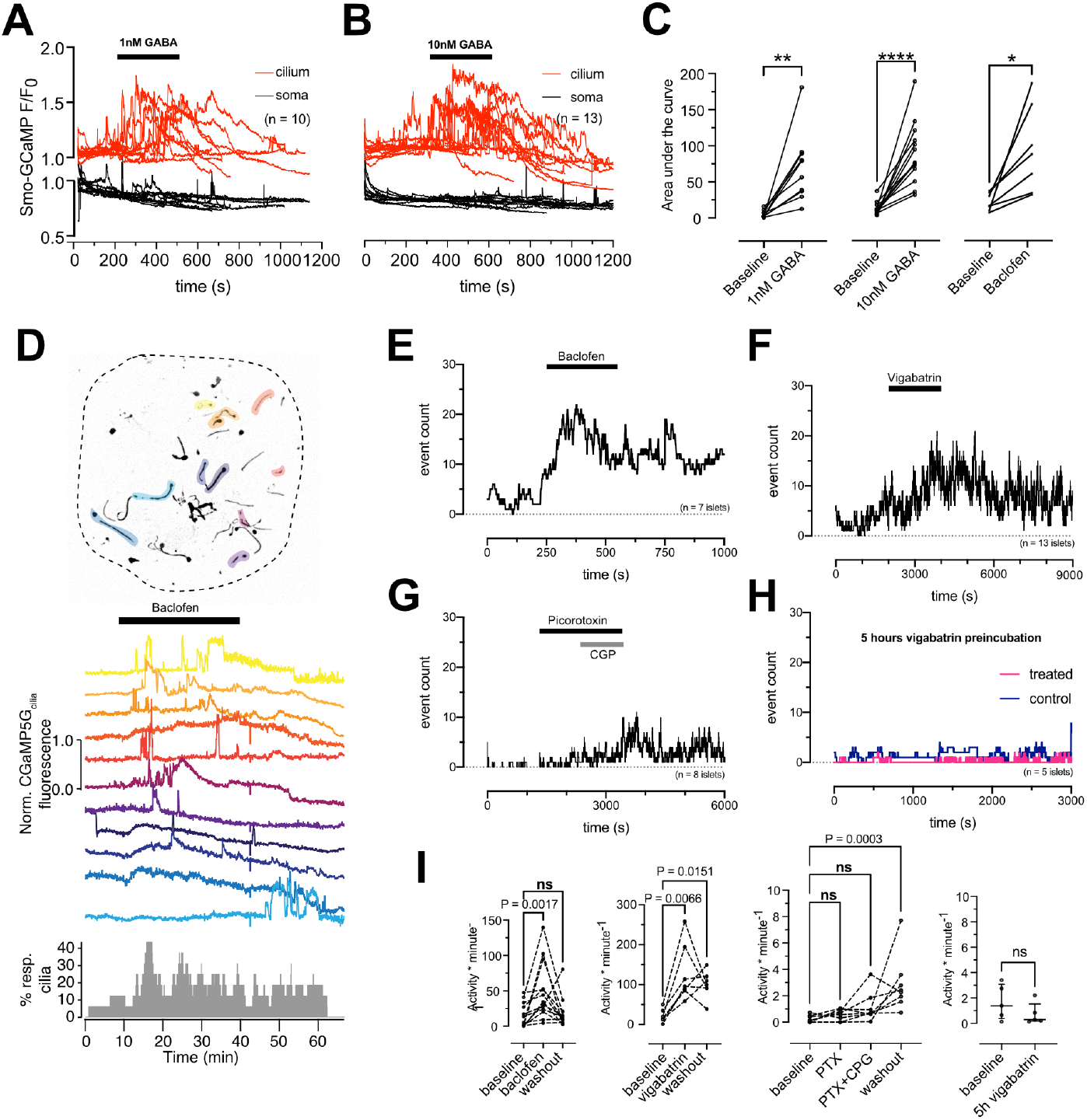
GABA triggers Ca^2+^ entry into primary cilia via GABA-B1 receptors. A. TIRF microscopy recordings of GCaMP5G fluorescence in the soma and cilium of MIN6 cells exposed to 1 nM GABA. Notice that the addition of GABA induces a rise of cilia Ca^2+^ without affecting the Ca^2+^ concentration in the soma. B. TIRF microscopy recordings of GCaMP5G fluorescence in the soma and cilium of MIN6 cells exposed to 10 nM GABA. Notice that the addition of GABA induces a rise of cilia Ca^2+^ without affecting the Ca^2+^ concentration in the soma. C. GABA (1 and 10 nM) and the GABA-B receptor agonist Baclofen induces increases in the cilia Ca^2+^ concentration in MIN6 cells (n=10 cells for 1 nM GABA; n=13 cells for 10 nM GABA; n=7 cells for Baclofen; * P<0.05; ** P<0.01; *** P<0.001; Student’s paired t-test). D. TIRF microscopy recordings of cilia Ca^2+^ concentration changes in intact mouse islets expressing Smo-GCaMP5G-Ch. Picture on top shows the distribution of cilia within the islet, with cilia exhibiting at least one spontaneous Ca^2+^ flash during the recording time highlighted in color. In the middle are shown Ca^2+^ recordings from the individual cilia within the islet during addition of the GABA-B receptor agonist Baclofen. The histogram at the bottom shows the overall cilia Ca^2+^ activity of the whole islet. E. Histogram showing the number of cilia Ca^2+^ flashes following the addition of baclofen (n=7 islets). F. Histogram showing the number of cilia Ca^2+^ flashes following the addition of vigabatrin (n=13 islets). G. Histogram showing the number of cilia Ca^2+^ flashes following the addition and washout of picorotoxin and CGP-35348 (n=8 islets). H. Histogram showing the number of cilia Ca^2+^ flashes following preincubation with DMSO (control) and vigabatrin for 5 h (n=5 islets for both conditions). I. Quantifications of the effect of baclofen, vigabatrin, picorotoxin+CGP-35348 and long-term vigabatrin treatment on cilia Ca^2+^ flashes. Statistical significance was assessed using the Friedman test followed by multiple comparisons.

**Figure 7.**
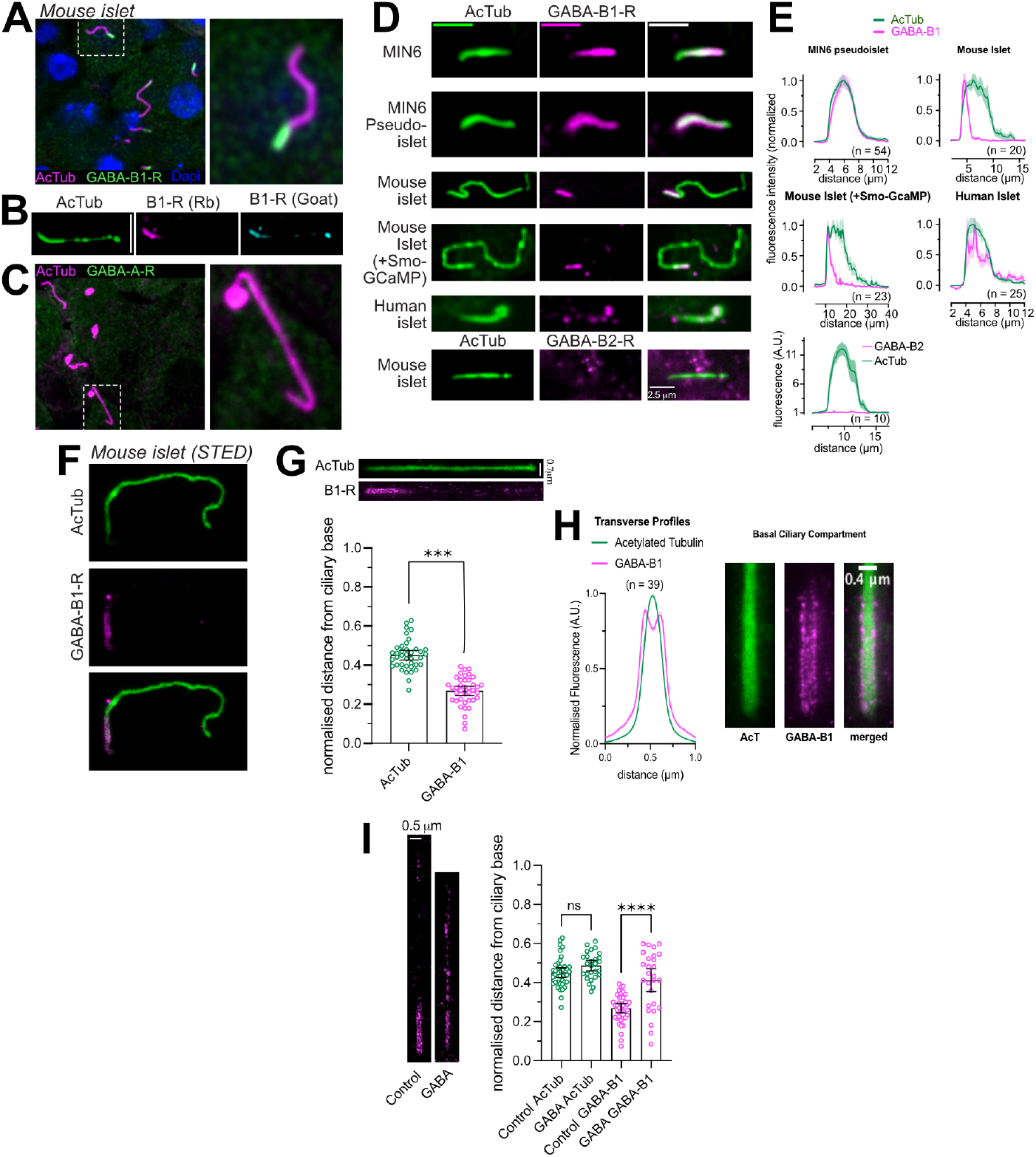
GABA-B1 receptors localize to the cilium and are mobilized by agonist binding. A. Immunofluorescence staining of a mouse islet showing acetylated tubulin in magenta, GABA-B1 receptors in green and the nuclei in blue (Dapi). B. Immunofluorescence staining of a mouse islet showing acetylated tubulin in green and GABA-B1 receptors in blue (anti-goat) and magenta (anti-rabbit). C. Immunofluorescence staining of a mouse islet showing acetylated tubulin in magenta and GABA-A receptors in green. D. Representative images showing the cilia localization of GABA-B1 receptors (magenta) to the primary cilium (green) in MIN6 cells, MIN6 pseudo-islets, mouse islet cells, mouse islet cells transduced with Ad-Smo-GCaMP5G-mCh and human islet cells. At the bottom is shown a cilium from a mouse islet stained for acetylated tubulin (green) and GABA-B2 receptors (magenta). E. Quantifications of the cilia distribution of GABA-B1 receptors in the different cell preparations. F. STED images of a mouse islet cell cilia stained for acetylated tubulin (green) and GABA-B1 receptors (magenta). G. Center-of-mass of acetylated tubulin and GABA-B1 receptors from “straightened” cilia (example above) of mouse islets (n=39, cilia from 25 islets). H. Distribution of GABA-B1 receptors at the base of the cilium quantified from STED images. Notice that the receptors localize peripheral to the microtubule core of the cilium (means ± S.E.M. for 39 cilia from 25 islets). I. Center-of-mass analysis of mouse islet cilia stained for acetylated tubulin and GABA-B1 receptors under control conditions and following a 10 min incubation with 10 nM GABA. Notice that the addition of GABA results in a shift in the center of mass towards more distal regions of the cilium (n=39, control and n=27, GABA cilia from at least 25 islets each, **** P<0.0001, One Way ANOVA with Multiple Comparisons).

## DISCUSSION

Functional studies of the mammalian primary cilium have primarily been carried out in cell lines, with the exception of pioneering work conducted with isolated olfactory neurons ^33,34^. Here we employed the islet of Langerhans as a model system that preserves organ like features and allows exploration of ciliary signaling by imaging approaches with very high resolution. Two features of this preparation are of crucial relevance in this regard; cells are arrested in the G_0_ stage of cell cycle (which means that their cilia completely develop a specialized phenotype) and a variety of endogenously produced signaling molecules are still operative in the isolated islet (which implies that ciliary pathways could be activated by signals also present *in vivo*). This is exemplified by the fact that both endogenous and overexpressed Smoothened shows a constitutive ciliary localization in islet cells, likely due to the endogenous production and release of Hedgehog ^35,36^. The use of this model benefitted our investigation of ciliary functionality by providing evidence of an efficient isolation of the cilium against changes in the cytosolic Ca^2+^ concentration, and enabled the identification of a GABA/GABA-B1 receptor signaling pathways that is coupled to cilia Ca^2+^ signaling. To the best of our knowledge, this is also the first example where primary cilia are shown to exhibit a clear, receptor-dependent, antenna-like sensing function within an intact mammalian organ.

A main finding in this study is that the primary cilium of islet cells is isolated against changes in cytosolic Ca^2+^. This is different from previous work on epithelial cells, where Ca^2+^ was shown to freely diffuse between the cytosol and cilium ^5,6^. The reason for this discrepancy is not clear, but one possibility is that efficient extrusion is more easily accomplished in the very long cilia of islet cells. It is also possible that this mechanism is more important in excitable cells, where regular changes in the cytosolic Ca^2+^ concentration are an essential part of normal cell function. Such an isolation is therefore a fundamental requirement if Ca^2+^ is to function as a cilia-intrinsic second messenger generated downstream of cilia-localized receptors and ion channels. Seclusion of the cilium is achieved by potent Ca^2+^ extrusion concentrated at the base of the cilium. The finding that amiloride is able to slow down clearance kinetics of both spontaneously generated Ca^2+^ signals and elevations induced by photorelease of caged Ca^2+^ points to a key role of the Na^+^/Ca^2+^ exchanger. However, amiloride is a rather non-selective inhibitor and other mechanisms of Ca^2+^ extrusion may also be involved. We also show that when β-cells are exposed to elevated glucose concentrations, the ensuing cytosolic Ca^2+^ oscillations are followed by a corresponding phase-locked, antiparallel fluctuation in the cilium, and this coupling is lost whenever the mechanisms generating the cytosolic activity are perturbed, implying a role of the membrane potential in ciliary Ca^2+^ extrusion. A similar mechanism of isolation has been observed in olfactory neurons, which are specialized, multi-ciliated cells ^33^. An interesting analogy can also be drawn between the primary cilium and the dendritic spine, another specialized structure with a unique Ca^2+^ signature and dimensions that resemble the cilium (1 μm long, 100 nm in diameter). Ca^2+^ changes in dendritic spines do not propagate into the dendrite, largely due to efficient extrusion ^37,38^. Ca^2+^ buffering may also help in shaping the cilia Ca^2+^ signals. Recent proteomic analysis of primary cilia has identified numerous Ca^2+^ -binding proteins that may be important for local Ca^2+^ buffering within the cilium ^39^. The generation of localized Ca^2+^ concentration increases in the cilium that failed to propagate throughout the structure implies a combination of efficient extrusion and local buffering. It would be relevant to determine whether these microdomains are related to the recently described actin corrals that constitute functional subunits of the cilium ^40^ or if they reflect the distinct localization of ion channels or Ca^2+^ binding proteins that might prevent unrestricted spreading of Ca^2+^ along the cilium.

Activation of muscarinic receptors caused a dramatic increase of Ca^2+^ at the base of the cilium, raising the possibility of an apposition with the ER, which is the main intracellular Ca^2+^ store. Although proximity between the ER and the cilia base has not been demonstrated, the ER plays a fundamental role in generating similar Ca^2+^ microdomains at the immunological synapse, a structure that bears many similarities to the primary cilium ^41,42^. An alternative possibility is that IP_3_ generated downstream of the muscarinic receptors instead mobilize Ca^2+^ from the Golgi apparatus ^43^, an organelle involved in both ciliogenesis and delivery of cargo to the mature cilium ^44^. The relevance of the effective isolation of the cilium emerges from the fact that Ca^2+^ is a key second messenger in this organelle and the occurrence of spontaneous Ca^2+^ flashes endorse this concept. The possibility of close proximity between the cilium and ER or Golgi allows us to speculate that ciliary Ca^2+^ transients could convey information to the cytoplasm by a yet to be described amplification performed by Ca^2+^ induced Ca^2+^ release. Indeed, we do occasionally observe a small, spatially restricted, rise in cytosolic Ca^2+^ as a Ca^2+^ wave propagates through the cilium and reaches the base (see Fig. 5A, B and D). Interestingly, similar bidirectional communication has been proposed to occur between the cilium and cytosol in olfactory neurons through a mechanism involving cilia-proximal mitochondria ^45^.

GABA is released from pancreatic β-cells in a glucose-independent manner, likely involving plasma membrane localized transporters ^18^. GABA exerts multiple effects on β-cells, including both acute modulation of insulin secretion and more long-term effects, such as proliferation and cell-fate maintenance ^16,17^. Most of these effects are thought to emanate downstream of ionotropic GABA-A receptors, which seem to be the dominating GABA receptor in the cell body ^17^. We now show that islet β-cells also express GABA-B receptors, and that the localization of these receptors is restricted to the primary cilium. This very distinct localization to, and action in, primary cilia could easily have been overlooked in earlier studies. We observe that the GABA-B1 receptors are enriched at the base of the cilium, which is at odds with the fact that the GABA-induced Ca^2+^ signalling typically originated from the medial to distal regions of the cilium. One explanation could be that the proximal accumulation of GABA-B1 represents a population of receptors leaving the organelle after activation, as has been described for other cilia receptors ^44^. Alternatively, that segment may be a functional representation of the inversin compartment ^32^, where the receptors are trapped in an inactive state. Agonist binding could relieve the inhibition, enabling diffusion to more distal segments where they could interact with signalling partners. Consistent with this notion, we observe receptor mobilization in response to shortterm incubation with GABA. How GABA binding to GABA-B1 receptors on the cilia surface leads to an increase in cilia Ca^2+^ is not clear, but most likely involves a non-canonical pathway that does not involve the GABA-B2 receptor subunit. This is supported by the observations that we failed to detect GABA-B2 by immunostaining and that the GABA-B1/2 antagonist CGP-35348 failed to abolish spontaneous Ca^2+^ flashes ^46^. The lack of effect of nifedipine and verapamil suggests that Ca^2+^ influx into the cilium is not mediated by voltage gated Ca^2+^ channels, although GABA-B receptors signalling negatively regulate voltage-dependent Ca^2+^-influx in neurons in a G-protein-dependent manner ^47,48^. One possibility is that ciliary Ca^2+^ influx is mediated by members of the transient receptor potential family (TRP channels) since most of them are non-selective cation channels and many also localize to the primary cilium ^3^. However, the fact that mechanical stimulation did not elicit any Ca^2+^ change speaks against this argumentation, since many TRP channels are believed to respond to such stimulations. Another possibility is that the Ca^2+^ flashes are mediated by cyclic nucleotide-gated channels (CNG), as is the case in olfactory neurons ^33^. GABA is an extremely versatile neurotransmitter and during development it switches from being an excitatory signal into an inhibitory one, mostly because of the reversal of the Cl^-^ gradient across the plasma membrane making GABA-A mediated currents hyperpolarizing ^49^. But there are also studies demonstrating increases of Ca^2+^ currents following activation of GABA-B receptors ^50,51^. Perhaps, the activity we show here represents an evolutionary atavism of GABA-B receptors.

The biological significance of the GABA-mediated ciliary Ca^2+^ signalling remains to be determined. GABA is released in a glucose-independent manner from islet cells ^18^, but interestingly we note that the GABA response in cilia is suppressed under conditions that stimulate insulin secretion, such as high glucose concentrations and direct depolarization. This implies that the mechanism is temporally uncoupled from insulin secretion and most prominent under resting conditions. GABA has previously been demonstrated to affect cell fate properties in islets cells, and also shown to have the ability to affect both proliferation and differentiation of mature β-cells ^52,53^, although these findings have been questioned in more recent studies ^54,55^. Interestingly, the cilium is ascribed similar properties in many cell types ^56^. It is therefore tempting to speculate that the ciliary GABA pathway may crosstalk with other cilia signalling pathways, such as Wnt and Hedgehog, either indirectly via second messenger generation or via sensitization of cilia receptors, similar to the role of GABA-B receptors in neurons and astrocytes ^57–59^. Elucidating the mechanism and function of GABA action on primary cilia represents an important future research direction.

## ACKNOWLEDGEMENTS

We are grateful to Bryndis Birnir, Anders Tengholm, Erik Gylfe and all the members of the OI-H lab for important input and suggestions on this work. We also acknowledge the excellent help with STED imaging provided by the staff at the BioVis facility at Uppsala University. This work was supported by grants from the Swedish research council, the Novo-Nordisk foundation, the Swedish diabetes foundation, the Family Ernfors foundation and Exodiab to OI-H.

