## Supplementary Material for "The β-cell primary cilium is an autonomous Ca^2+^ compartment for paracrine GABA signalling"

### SUPPLEMENTARY FIGURE LEGENDS

**Suppl. Figure 1. Characterization of a cilia-targeted $Ca^{2+}$ sensor.**

A, B. Principle for determining the enrichment of Smo-GCaMP5G-mCh within the primary cilium of islet cells imaged by TIRF microscopy. With a radius of the cilium of 100 nm and an evanescent wave penetration depth (d) of 80 nm, we estimate that a ratio of cilia-to-plasma membrane fluorescence >1.288 indicate cilia enrichment.

C. Quantifications of fluorescence intensities of Smo-GCaMP5G-mCh from the plasma membrane and cilia of islet cells. The average cilia-to-plasma membrane fluorescence ratio is 2, which indicate strong cilia enrichment (n=113 soma and 95 cilia; Student's 2-tailed unpaired t-test).

D. Immunofluorescence staining of a MIN6 cells expressing Smo-GCaMP5G-mCherry using antibodies against acetylated tubulin (green) and mCherry (magenta).

E. Test of the topology of Smoothened in the cilia membrane using a version where the pH-sensitive green fluorescence protein pHluorin is tagged at the N-terminus (extracellular). MIN6 cells expressing ss-pHluorin-Smo were imaged on the stage of a TIRF microscope and exposed to a weakly acidic buffer (pH 6) to quench surface pHluorin fluorescence (means ± S.E.M. for 11 cells form 3 experiments).

F. Test of the topology of ss-pHluorin-Smo by the addition of proteinase K while monitoring pHluorin fluorescence from the cilia of MIN6 cells by TIRF microscopy (means ± S.E.M. for 9 cells from 3 experiments).

G. Dose-response curves for Smo-GCaMP5G-mCh fluorescence changes in the plasma membrane (green) and cilia (black) of permeabilized MIN6 cells (means ± S.E.M. for 8 cells from 3 experiments).

H. Measurements of cilia length in islets cells from mouse pancreatic sections (green), isolated mouse islets (red) and mouse islets transduced with Smo-GCaMP5G-mCh for 48 h (black).

I. TIRF microscopy images of a MIN6 cell expressing Smo-GCaMP5G-mCh and loaded with caged $Ca^{2+}$. White dot marks site of local UV-induced photolysis of the caged $Ca^{2+}$. Graph below shows the GCaMP5G fluorescence increase in the cell body (red) and cilium (black) following uncaging (means ± S.E.M., n=9 cells).

J. Scatter plots showing the lack of correlation between the expression level of Smo-GCaMP5G-mCh and half-life of $Ca^{2+}$ decay in the cilium (left) and cell bodies (right) of MIN6 cells following photolysis of caged $Ca^{2+}$ (n=24 cells).

K. TIRF microscopy images showing lack of recovery of mCherry fluorescence following local photobleaching (dashed white circle) of MIN6 cells expressing Smo-GCaMP5G-mCh). Graphs to the left shows the recovery of fluorescence after photobleaching in the plasma membrane (top) and cilium (bottom) (means ± S.E.M.; n=14 cells).

L. TIRF microscopy images of a MIN6 cell expressing the cilia-targeted pH-sensor $5HT_6$-Venus-CFP under resting conditions. Notice that the venus/CFP ratio is higher in the cilium.

M. Means ± S.E.M. (n=30 cells) of line-profiles drawn from cilia to cell body across the base of the cilium in TIRF microscopy images of MIN6 cells expressing $5HT_6$-Venus-CFP. Notice that the venus/CFP ratio is higher in the cilium, indicating more alkaline pH.

N. Cilia and soma venus/CFP fluorescence ratios in MIN6 cells expressing $5HT_6$-Venus-CFP (n=30 cells, Student's 2-tailed paired t-test).

**Suppl. Figure 2. $Ca^{2+}$-independent changes in GCaMP5G fluorescence induced by 20 mM glucose.**

A. Ratiometric recording (black) of GCaMP5G (green) and mCherry (magenta) fluorescence from a mouse islet cell cilium expressing Smo-GCaMP5G-mCh. Notice that glucose causes an immediate increase in GCaMP5G fluorescence but is without effect on mCherry fluorescence.

B. Ratiometric (GCaMP5G/mCherry) recording from the cilium of a mouse islet cell exposed to 11 mM glucose followed by the addition of hyperpolarizing diazoxide (250 µM) and carbachol (10 µM). Notice that the addition of diazoxide suppresses the glucose-induced $Ca^{2+}$ oscillations but does not bring the GCaMP5G/mCh ration back to resting levels.

C. TIRF microscopy recording of GCaMP6 fluorescence from a transgenic mouse islet β-cell expressing GCaMP6 under control of the insulin promoter. The islet was exposed to a step increase in the surrounding glucose concentration, from 3 mM to 20 mM, followed by addition of the hyperpolarizing agent diazoxide. Notice that in contrast to Smo-GCaMP5G-mCh (see Suppl. Fig. 2A), glucose does not cause an immediate increase in GCaMP fluorescence, and the glucose-induced increase in GCaMP fluorescence is suppressed to resting levels in the presence of diazoxide.

D. Fluorescence changes from α-toxin-permeabilized MIN6 cells expressing Smo-GCaMP5G-mCh (green/dashed-black) or plasma membrane-anchored R-GECO ($Lyn_{11}$-R-GECO; magenta) following exposure to intracellular-like buffers containing the indicated $Ca^{2+}$ and ATP concentrations.

E. Quantifications of GCaMP5G and $Lyn_{11}$-R-GECO fluorescence changes in permeabilized cells exposed to the indicated intracellular buffers (n=13-16 cells, **$P<0.01$, 2-tailed paired Student's t-test).

F. Venus/CFP ratio changes in the cell body (red) and cilia (black) of MIN6 cells expressing cilia-targeted $5HT_6$-Venus-CFP and exposed to an increase in the surrounding glucose concentration from 3 mM to 20 mM (means ± S.E.M. of 21 cells).

G. Venus/CFP ratio changes in the cell body (red) and cilia (black) of MIN6 cells expressing cilia-targeted $5HT_6$-Venus-CFP and exposed to 20 mM $NH_4Cl$, which causes strong alkalinization of the cytosol (means ± S.E.M. of 21 cells).

**Suppl. Figure 3. Characterization of spontaneous $Ca^{2+}$ flashes in MIN6 cells and mouse islet cells.**

A. Spontaneous $Ca^{2+}$ "flash" in the primary cilium of a MIN6 cells expressing Smo-GCaMP5G-mCherry. Notice how the flash originates in the tip of the cilium and propagates towards the base and how this coincides with a small local rise of $Ca^{2+}$ in the cilia-adjacent cytosol.

B. Characteristics of cilia $Ca^{2+}$ flash diffusion rate, duration, amplitude and site of origin in MIN6 cells expressing Smo-GCaMP5G-mCh (n=22).

C. A spontaneous $Ca^{2+}$ flash in a mouse islet cell expressing Smo-GCaMP5G-mCh and kept in a buffer containing 20 mM glucose and 250 µM diazoxide. Notice how the wave propagates from tip to base and how the strength of the flash is diminished as it approaches the base.

D. A spontaneous $Ca^{2+}$ flash in a mouse islet cell expressing Smo-GCaMP5G-mCh and kept in a buffer containing 20 mM glucose, 250 µM diazoxide and amiloride (200 µM). Notice the more homogenous rise of $Ca^{2+}$ along the cilium and the longer duration of the flash.

E. Characteristics of cilia $Ca^{2+}$ flash diffusion rate, duration and amplitude in mouse islet cells expressing Smo-GCaMP5G-mCh and kept in 3 mM glucose, 20 mM glucose and diazoxide or 20 mM glucose, diazoxide and amiloride (* $P<0.05$, ** $P<0.01$, *** $P<0.001$). Diffusion rate: Kruskal-Wallis test with multiple comparisons, 3G n=37 cells from 30 islets; 20G n=6 cells from 4 islets; 20G+amiloride n=7 cells from 5 islets. Duration: Brown-Forsythe and Welch ANOVA test with multiple comparisons, 3G n=31 cells from 30 islets; 20G n=8 cells from 4 islets; 20G+amiloride n=10 cells from 5 islets. Amplitude: Kruskal-Wallis test with multiple comparisons, 3G n=33 cells from 30 islets; 20G n=8 cells from 4 islets; 20G+amiloride n=10 cells from 5 islets.

**Suppl. Figure 4. GABA-B1 receptor distribution in the cilium after stimulation.**

A, B. Quantifications of line profiles drawn along cilia of MIN6 cells immunostained for acetylated tubulin (A) and GABA-B1 receptors (B). The two lines are form cells treated for 48h with control siRNA (Control) or siRNA against the GABA-B1 receptor (GABA-B1-R KD). Notice how the GABA-B1 immunoreactivity is reduced in cells treated with GABA-B1 R siRNA.

C. Cilia fluorescence intensity from control and GABA-B1 receptor KD MIN6 cells immunostained for acetylated tubulin (left) or GABA-B1 receptors (right). Unpaired 2-tailed Student's t test, n=15 from both groups.

D. Quantifications of GABA-B1 receptor fluorescence intensity and distribution along primary cilia of mouse islet cells under control conditions (black) and following 10 min stimulation with 100 nM of the GABA-B receptor agonist Baclofen (red). Data presented as means ± S.E.M. for the indicated number of cells.

E. Quantifications of GABA-B1 receptor fluorescence intensity and distribution along primary cilia of mouse islet cells under control conditions (black) and following 10 min stimulation with 100 nM GABA (red). Data presented as means ± S.E.M. for the indicated number of cells.

F. GABA-B1 receptor distribution in mouse islet cilia under control conditions and following stimulation for 10 min 100 nM GABA or 100 nM Baclofen. Data presented are calculations of the area under the curve of line profiles drawn along cilia of mouse islet cells immunostained for acetylated tubulin and GABA-B1 receptors. Representative images of cilia from a control cell and a cell exposed to Baclofen are shown to the right (green, acetylated tubulin; magenta, GABA-B1 receptor). Anova with multiple comparisons; n=63 (control) and n=46 (baclofen); n=175 (control) and n=133 (GABA).

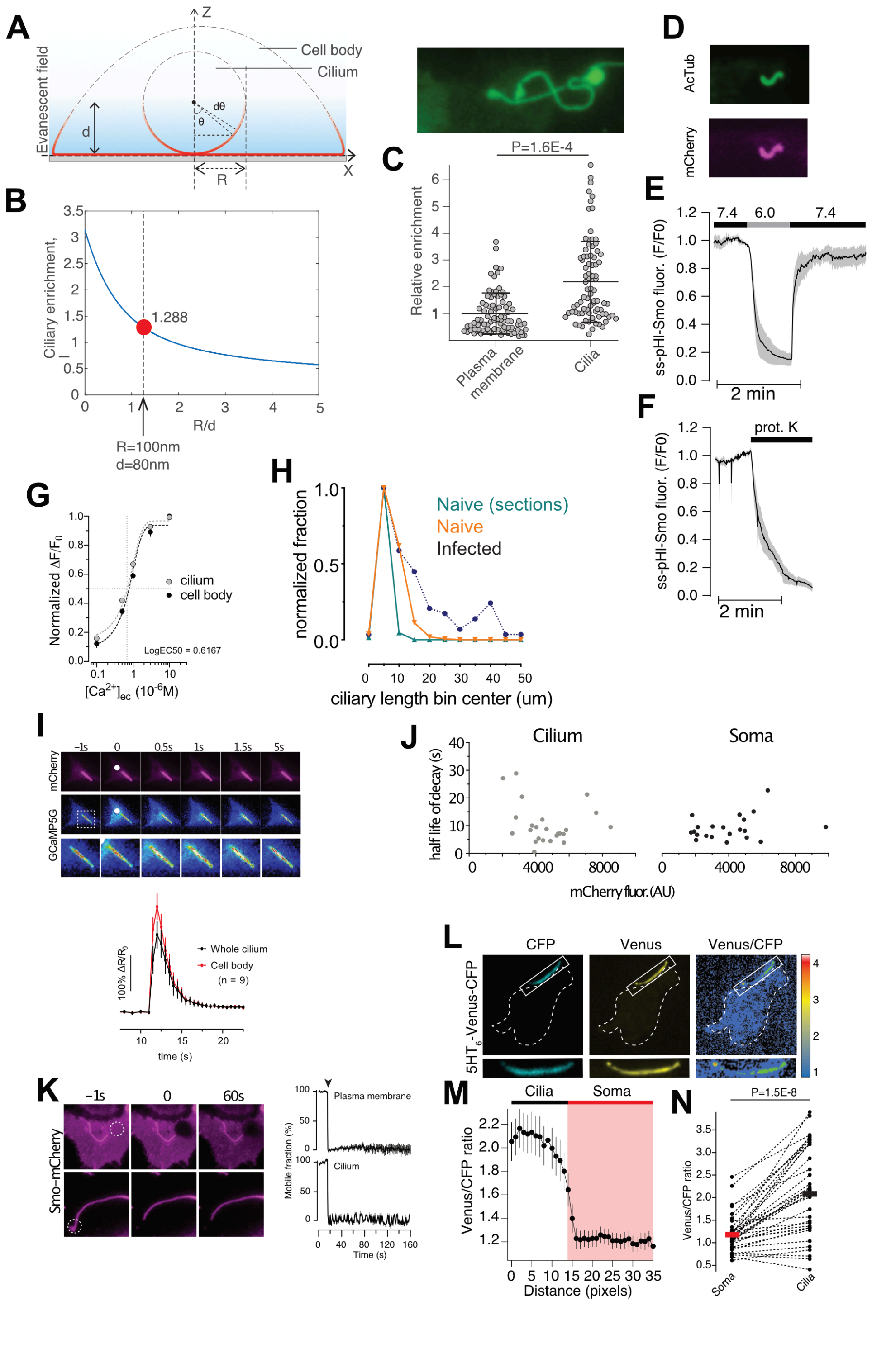


Suppl. figure 1
Sanchez et al

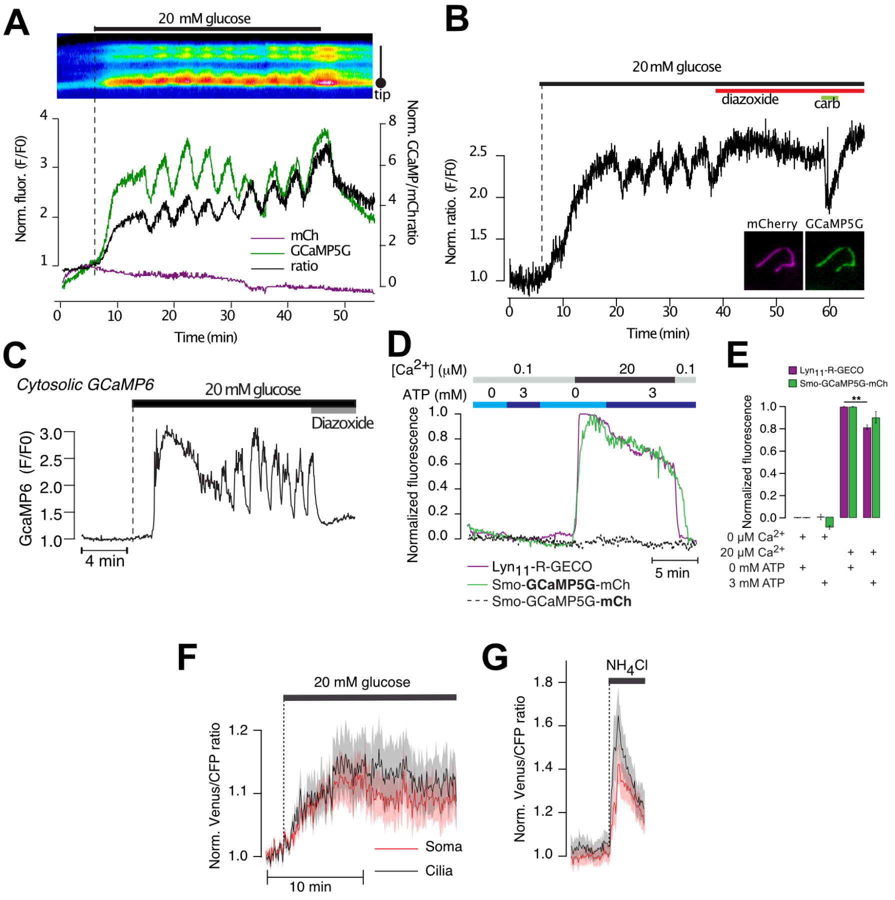


Suppl. figure 2
Sanchez et al

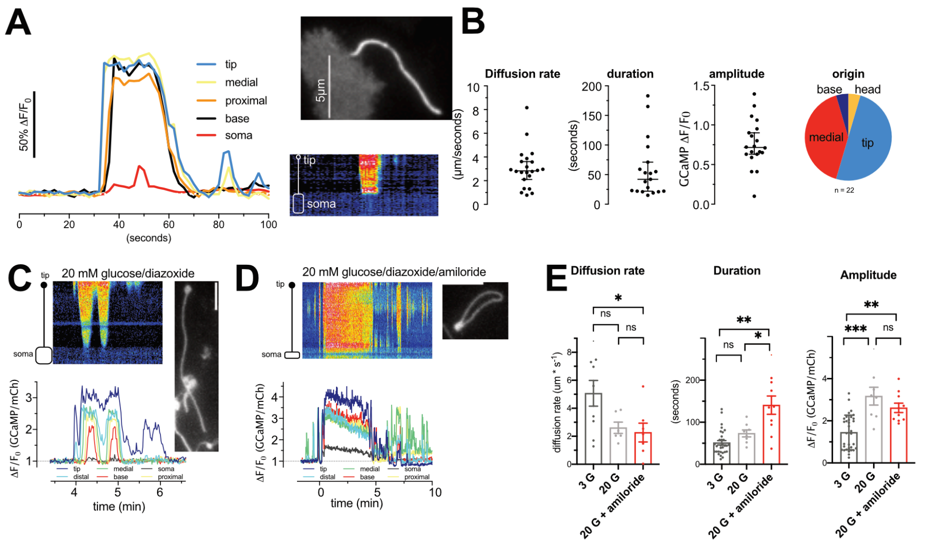


Suppl. figure 3
Sanchez et al

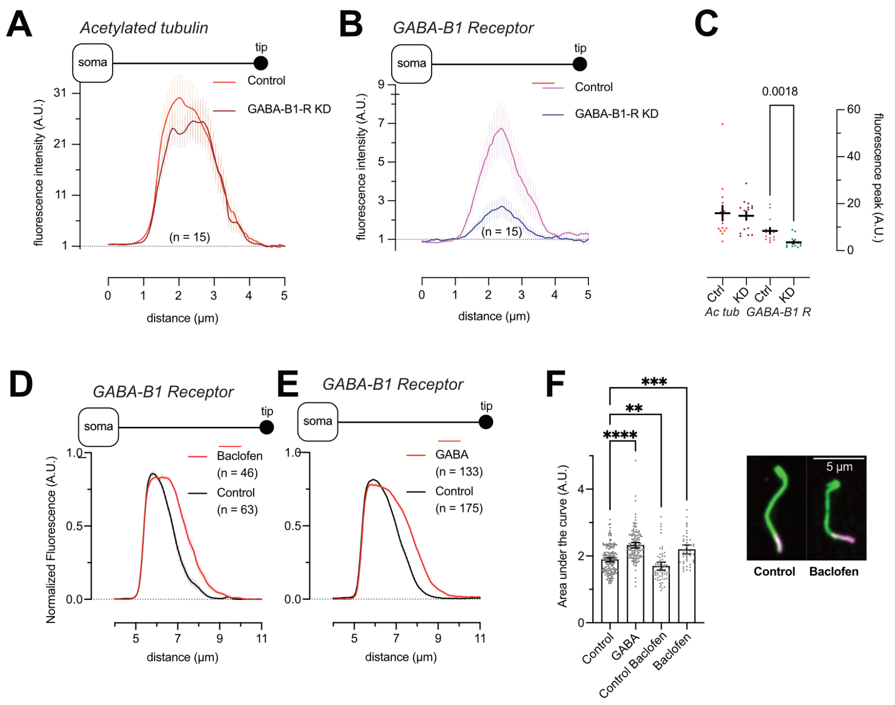


Suppl. figure 4
Sanchez et al
